# Genetic access to an obligate cyanobacterial endosymbiont within the shoot meristem of fern hosts

**DOI:** 10.64898/2026.02.17.706265

**Authors:** Cristina Sarasa-Buisán, Erbil Güngör, Enrique Flores, Peter Lindblad, Sandra Nierzwicki-Bauer, Henriette Schluepmann

## Abstract

Eukaryote-associated microbes are ubiquitous, but their essential roles in the development and ecology of their host is yet to be fully understood, partly because complex associations cannot be reconstituted and, in many instances, the genetic tools to elucidate those roles are not available. Here, we report the conjugative transfer of DNA into *Nostoc azollae* within two *Azolla* fern hosts. *N. azollae* is a filamentous, N_2_-fixing, heterocyst-forming cyanobacterium which is an obligate endosymbiont of the complex microbial community associated with the floating ferns of the genus *Azolla*. The cyanobiont provides fixed nitrogen to its host, supporting maximum growth rates without any N-fertilizer and making *Azolla* symbioses both ecologically and agriculturally important. Triparental mating protocols and fluorescent reporter detection were optimized for the cyanobiont isolated from the fern, allowing further demonstration of heterologous gene expression in *N. azollae* driven by several promoters, including some of a CRISPR-associated transposon (CAST) system. *Azolla* was then treated with a cytokinin hormone to render fern shoot apices amenable to *in planta* conjugation, permitting DNA transfer to, and stable gene expression in two distinct developmental stages of *N. azollae* within *Azolla*. These included (i) cells of filaments from the Shoot Apical *Nostoc* colony, the only cyanobacterial stem-cell population vertically transmitted across fern generations, and (ii) cells from differentiated filaments in early formed *Azolla* leaf cavities. Our approach represents a technically groundbreaking advance for the genetic engineering of cyanobacterial endosymbioses that may be useful for other symbiotic systems, opening a pathway to investigate these important biological entities.

## Introduction

Ferns from the genus *Azolla* and their cyanobiont *Nostoc azollae* form one of the most intimate endophytic symbioses among plant–cyanobacterial associations (1, 2). *N. azollae* is a filamentous, N_2_-fixing cyanobacterium that transfers to its host about 40% of the N it fixes in heterocysts (3, 4); it is a vertically inherited obligate symbiont that exhibits extensive genome erosion (2, 5). *N. azollae* is supported at least partially by sugar received from its host (6, 7). This mutualism enables *Azolla* to thrive in nitrogen-poor environments with traditional uses as a biofertilizer in rice crops. Today, *Azolla* cultivation is also promising for a high-protein food and feed, a biofuel, bioremediation, greenhouse gas mitigation, and pest control (8–14). However, its adoption will require that biomass yields are stabilized, which depends in part on the development of strains that are resistant to common biotic stressors.

The *Azolla*–*Nostoc* symbiosis remains genetically intractable. On the plant side, progress has been made through protoplast induction, sexual crossing, cryopreservation, (meta)genome sequencing and transcriptomic analyses (15–17), but stable transformation of *Azolla* has yet to be achieved. On the cyanobiont side, the obligate endosymbiotic nature of *N. azollae* restricts experimental approaches to *in planta* DNA delivery methods. Moreover, the *N. azollae* genome has lost most genes involved in natural competence (See Table S1 from Wendt & Pakrasi 2019(18)), limiting methods of DNA delivery to active DNA mobilization such as bacterial conjugation. Therefore, the question arises, after about 100-Myr of evolution of *N. azollae* within *Azolla* (19, 20), can *N. azollae* still undergo conjugation?

In this work we assess whether an *Escherichia coli*-mediated conjugation with an eYFP- monitoring system, optimized in model cyanobacteria, could be adapted to *N. azollae* firstly outside the fern and then *in planta*. Targeted *N. azollae* reside in the host shoot apex, where a colony of filaments is maintained that provides inoculum for the developing leaves, and also in reproductive organs of the fern fronds. Additionally, we show the delivery of a large genome editing tool, the CRISPR-associated transposon (CAST) system. CAST is a powerful tool, enabling precise, large DNA insertions independent from host repair pathways (21), which has been validated in a model free-living, heterocyst-forming cyanobacterium, *Anabaena* sp. PCC 7120 (hereafter *Anabaena*), for YFP-tagged deletions (22).

## Results

### Tracing early conjugation events in cyanobacteria

To assess the feasibility of conjugative DNA transfer in the obligate cyanobiont *N. azollae*, we developed a fluorescence-based methodology to monitor early DNA transfer events in cyanobacteria using the *Anabaena* model. Additionally, we studied whether early conjugation events could be detected using, as cargo, a plasmid carrying the sgRNA-guided transposition CAST system (22). To enable reliable monitoring of conjugation events, we constructed an RSF1010-based plasmid containing a fluorescence reporter (enhanced Yellow Fluorescent Protein [eYFP]) driven by the strong synthetic *E. coli* promoter P_*trc*1O_ (23, 24). The resulting fluorescent cassette was tested either as an independent construct within the replicative backbone pCAT.000 (generating plasmid pL0) or by replacing the *P*_*rnpB*_*–*eYFP cassette in a previously described CAST plasmid (22) yielding updated versions of CAST plasmids (referred to as pCAST01 [Em^R^] and pCAST02 [Sp^R^Sm^R^]) ready to clone any sgRNA by one single Golden Gate reaction with *LguI* (Table S1).

Conjugation events of *Anabaena* receiving pL0 and pCAST02 targeting the *nifK* gene, hereafter referred as pCAST-7120 (Fig. 1A), were monitored 21- and 48-hours post-conjugation, as well as after 3 days of antibiotic selection. A scrape of cells from the conjugation plate (Fig. 1B) was transferred to a plate of BG11_0_ medium (which lacks combined nitrogen) and visualized using confocal microscopy. Conjugation events were detected in *Anabaena* as early as 48 hours post-mating, as eYFP signal localized to the cytoplasm of the exconjugants with either pL0 (Fig. 1C) or pCAST-7120 (Fig. 1D). At all observation time points, residual *E. coli* donor cells carrying pL0 or pCAST-7120 were also observed as ∼1-µm eYFP-expressing rods or as chain-like aggregates (characteristic of biofilms (25)), which were readily distinguishable from cyanobacterial cells. Conjugation events were verified by Lambda scans (500-700 nm), in which the fluorescence spectra of *Anabaena* exconjugants and controls were analyzed (Fig. 1E). A characteristic emission peak at 525 nm in both *E. coli* donor cells and cyanobacterial exconjugants confirmed the presence of the eYFP. Additionally, the scans verified that *Anabaena* cells displayed their typical autofluorescence derived from cyanobacterial pigments. This procedure was critical to distinguish true eYFP fluorescence from background cyanobacterial autofluorescence and *Azolla* tissue fluorescence.

**Figure 1.**
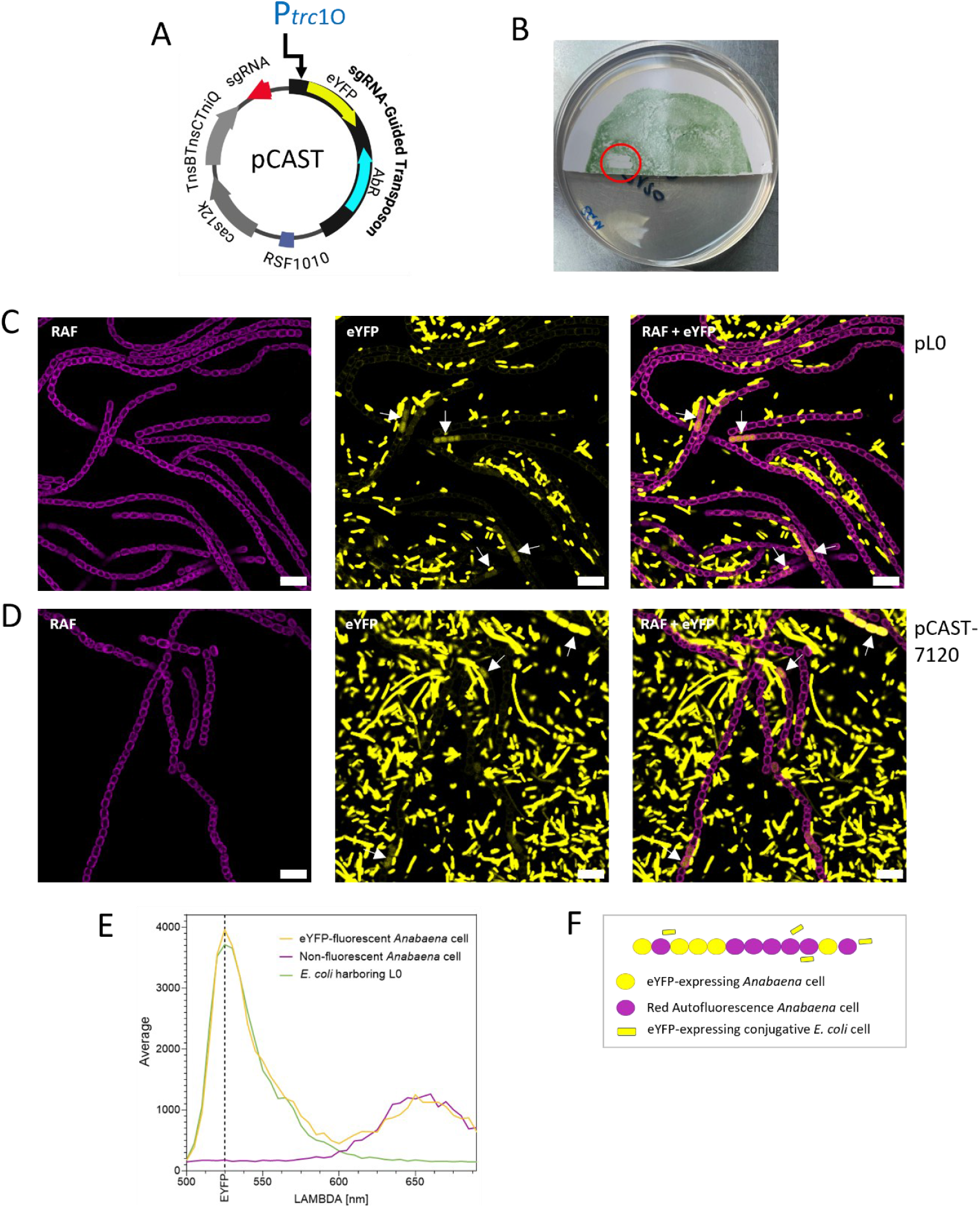
Tracing early conjugation events in *Anabaena* sp. PCC 7120 by confocal microscopy. (A) Schematic representation of the CAST plasmid pCAST-7120, featuring the P*trc*1O promoter for eYFP expression. (B) Conjugation plate of *Anabaena* 48 hours after conjugation highlighting in red the scrap taken 21 h after conjugation. Part of the conjugation plate was scraped up and re-streaked in a BG110 medium agar plate for visualization. (C,D) Confocal microscopy images displaying conjugation events at day 2 after conjugation with the control plasmid pL0 (C) and the pCAST-7120 plasmid including a sgRNA targeting *nifK* (D). Some conjugation events are marked with a white arrow. Note the presence of conjugative *E. coli* cells harboring cargo plasmids and expressing the eYFP. The conjugation (triparental mating) was performed as described in Materials and Methods. Red Auto Fluorescence (RAF, shown in magenta); enhanced Yellow Fluorescent Protein (eYFP). Images are representative of at least 5 independent experiments. Scale bar, 10 μm. (E) Example of lambda scan determination of specific eYFP signal in *E. coli* donor cells and *Anabaena* exconjugants. (F) Scheme depicting the different cells observed, expressing or not expressing the eYFP.

A spatial pattern of *Anabaena* exconjugant cells at early stages was observed and suggests that paired adjacent fluorescent cells may represent a single conjugation event followed by cell division, whereas non-contiguous fluorescent cells within the same filament are likely to represent multiple independent conjugation events in that filament (Fig. 1 and Fig. S1). Therefore, this criterion was used for subsequent quantifications of conjugation events. Noteworthy, several non-contiguous exconjugant cyanobacterial cells could display different levels of eYFP fluorescence intensity with either pL0 and pCAST-7120 (Fig. 1 and Fig. S1). This may result from variations in plasmid copy number, as has been observed previously with replicative plasmids in filamentous cyanobacteria (26), or, in the case of pCAST-7120, from the sum of eYFP expression from the replicative plasmid and transposon insertion events into the chromosome that may have already taken place. Because low fluorescence intensity may hinder early detection, this implies that observed conjugation events may be underestimated. Consequently, it is possible that conjugation had already occurred at earlier time points but remained undetectable at 21 hours. Modifications in the standard *Anabaena* conjugation protocol (Materials and Methods), such as incubation during the first 24 hours in darkness, extending the *E. coli* mating time (prior to mixing with *Anabaena*) from 2 to 4 hours, or the use of the Em^R^ pCAST01 version as the backbone (see pCSCSBT22 in Table S1), were tested to explore conditions that might be useful for subsequent *in planta* applications. However, these conditions did not result in an earlier detection.

### Isolated *Nostoc azollae* filaments can be conjugated *ex planta*

A conjugation protocol “*ex planta*” was established using *N. azollae* filaments extracted from fronds of *Azolla filiculoides* strain Galgenwaard (5) (Fig. S2), a sample called hereafter *Azolla* juice (7). The viability for several days of *N. azollae* filaments present in the *Azolla* juice was demonstrated previously (7). An adapted triparental mating protocol (Fig. S3A) was used with *E. coli* donor strains carrying either the control plasmid pL0 or the pCAST02-based pCAST-Nazo plasmids (pCAST- Nazo23, encoding an sgRNA targeting *nifK*; pCAST-Nazo24, encoding an sgRNA targeting the non-functional pseudogene *narB*) (Table S1 and Table S2). This yielded detectable conjugation events in *N. azollae* (Fig. 2 and Fig. S4, respectively), with eYFP fluorescence signals matching those of *E. coli* controls in Lambda scan analyses (Fig. 2C). Overall, exconjugants were detected within a time frame of 2-5 days after mating, which corresponds with the timing observed for the *Anabaena* control (Fig. 1).

**Figure 2.**
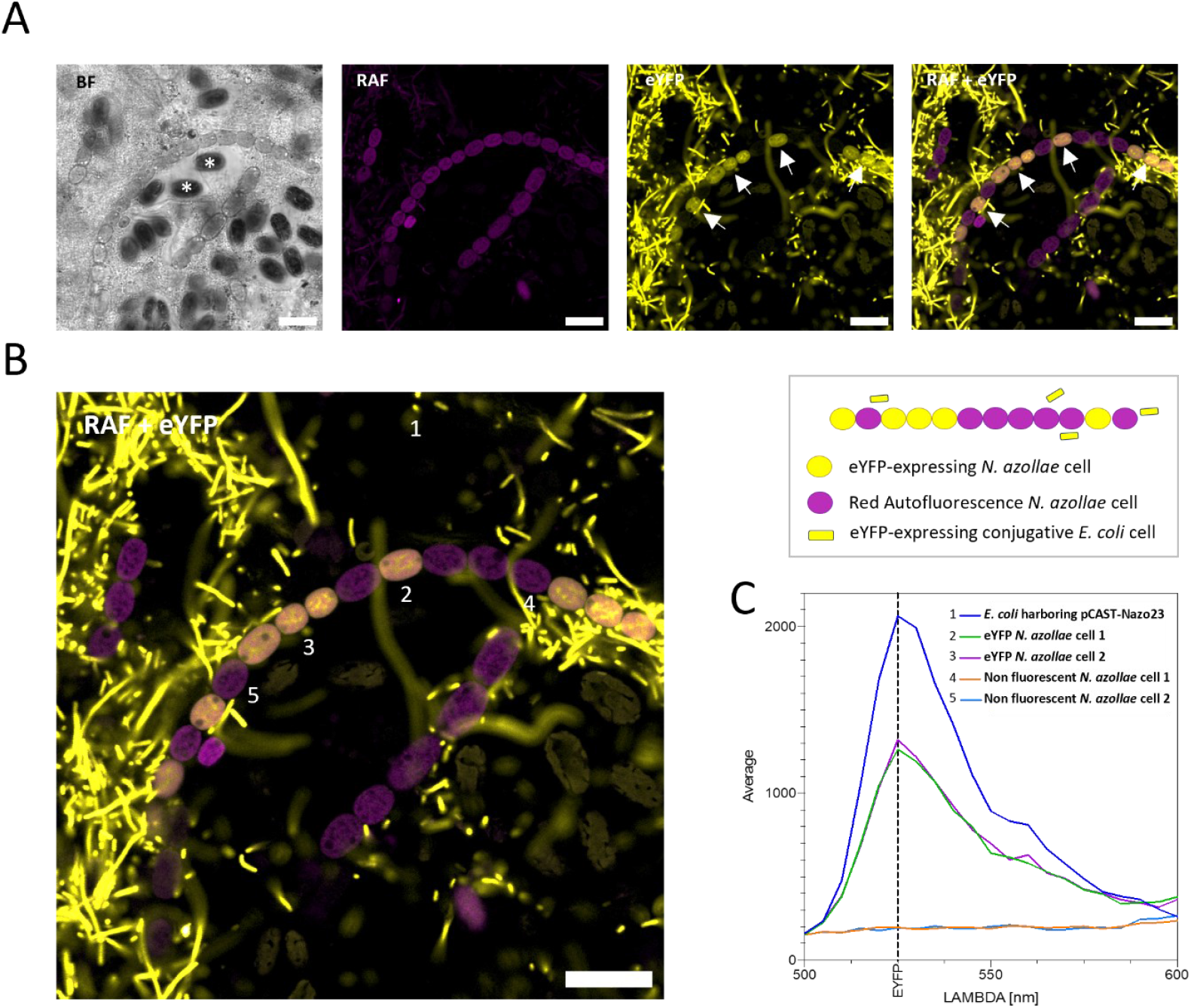
Conjugation events detected through eYFP expression in *Nostoc azollae* isolated from *Azolla filiculoides*. Conjugation of *N. azollae ce*lls extracted from *Azolla filiculoides* Galgenwaard was performed and analyzed as described in Materials and Methods. (A) Confocal imaging of *N. azollae* conjugation events in four detection channels: BF (brightfield), RAF (red autofluorescence, shown in magenta), eYFP, and merged eYFP + RAF. Several conjugation events of *N. azollae* cells are marked with a white arrow. Possible Eukaryotic algae, which were likely present on the surface of the fern, are shown with (*). (B) Enlarged view of the RAF+eYFP fluorescence channel. (C) Average fluorescence emission spectra corresponding to the numered cells in panel B. Note fluorescent eYFP *N. azollae* cells showing the same characteristic eYFP emission peak as *E. coli* harboring the eYFP expressing plasmid pCAST-Nazo23. Scale bar, 15 µm. A scheme depicting the different cells observed, expressing or not expressing the eYFP, is also included.

In several cases, eYFP-positive cells could be detected within non-adjacent cells in a single filament, suggesting independent conjugation events. In addition, contiguous eYFP-expressing cells were frequently found, consistent with conjugation followed by cell division (Fig. 2B). In a few cases, eYFP-positive vegetative cells were detected in filaments containing heterocysts (Fig. S4), which reflects that the *Azolla* juice harbors filaments in different developmental states, from leaf pockets (presence of heterocyst and larger cells) or from meristematic regions (narrowed-cells and absence of heterocysts (7)). Nonetheless, overall, conjugation events were frequently detected in short filaments or paired cells (Fig. S5), consistent with filament fragmentation observed in the sample due to the mechanical manipulation of the extraction protocol (7) (Fig. S2). After day 5 following conjugation, extracted cyanobacterial filaments subjected to conjugation showed signs of reduced viability and reached a near-complete disintegration.

To assess whether this protocol could be used with another host species, we carried out the conjugation assay with *N. azollae* filaments isolated from an *Azolla* line considered to be very robust, *Azolla anzali* (17). The same plasmid constructs that had been validated in *A. filiculoides* strain Galgenwaard were used for *A. anzali*. In this case, eYFP-positive cells could also be detected (Fig. S6), showing that conjugation can be achieved in *N. azollae* from different *Azolla* host species.

Together, these findings indicate that *N. azollae*, from two independent *Azolla* host species, have retained the functional traits required to receive DNA by conjugation despite its long-term obligate symbiotic lifestyle. Moreover, the results show that plasmids carrying the RSF1010 origin of replication and a P_*trc*1O_-driven reporter are functional for heterologous gene expression in *N. azollae*, and that the ∼16 kb pCAST derived constructs can be replicated and maintained in *N. azollae*, which will be necessary for the development of CAST-based tools in this symbiont.

### *In planta* conjugation of *Nostoc azollae*

Genetic manipulation of *N. azollae* that is transmitted to the developing host organs including the leaves and reproductive structures (and therefore to the next plant generation) requires that conjugative *E. coli* targets filaments in the Shoot Apical *Nostoc* Colony (SANC), which resides within the Shoot Apical Meristem (SAM) of *Azolla* (see Fig. 1 in Dijkhuizen et al. 2021 (17)). The SANC can be considered an *N. azollae* “stem-cell” colony that colonizes newly formed leaves and reproductive organs as they form (1, 27) (Fig. 3A). The SAM is normally covered by leaves such that the SANC is inaccessible to *E. coli* (Fig. 8A). To access the SANC, it was necessary to expose the SAM, which we achieved by treatment with the cytokinin 6-benzylaminopurine (BAP, Fig. S7, S8). Initial experiments with *A. filiculoides* strain Galgenwaard showed that BAP treatment severely weakened the plants, often compromising short-term survival required for early monitoring. Therefore, *A. anzali* was examined as a more robust alternative, owing to its faster growth under laboratory conditions. SAM exposure was induced with BAP (Fig. S7B), and 24 hours of exposure to hormone was sufficient to elicit open apexes detectable after 10 days (shown in Fig. S8). Longer incubation times did not increase the number of open apexes, suggesting that the response saturates within 24 h. The closed SAM phenotype with outgrowing meristems was restored approximately 1-month post-treatment onwards (Fig. S7C).

**Figure 3.**
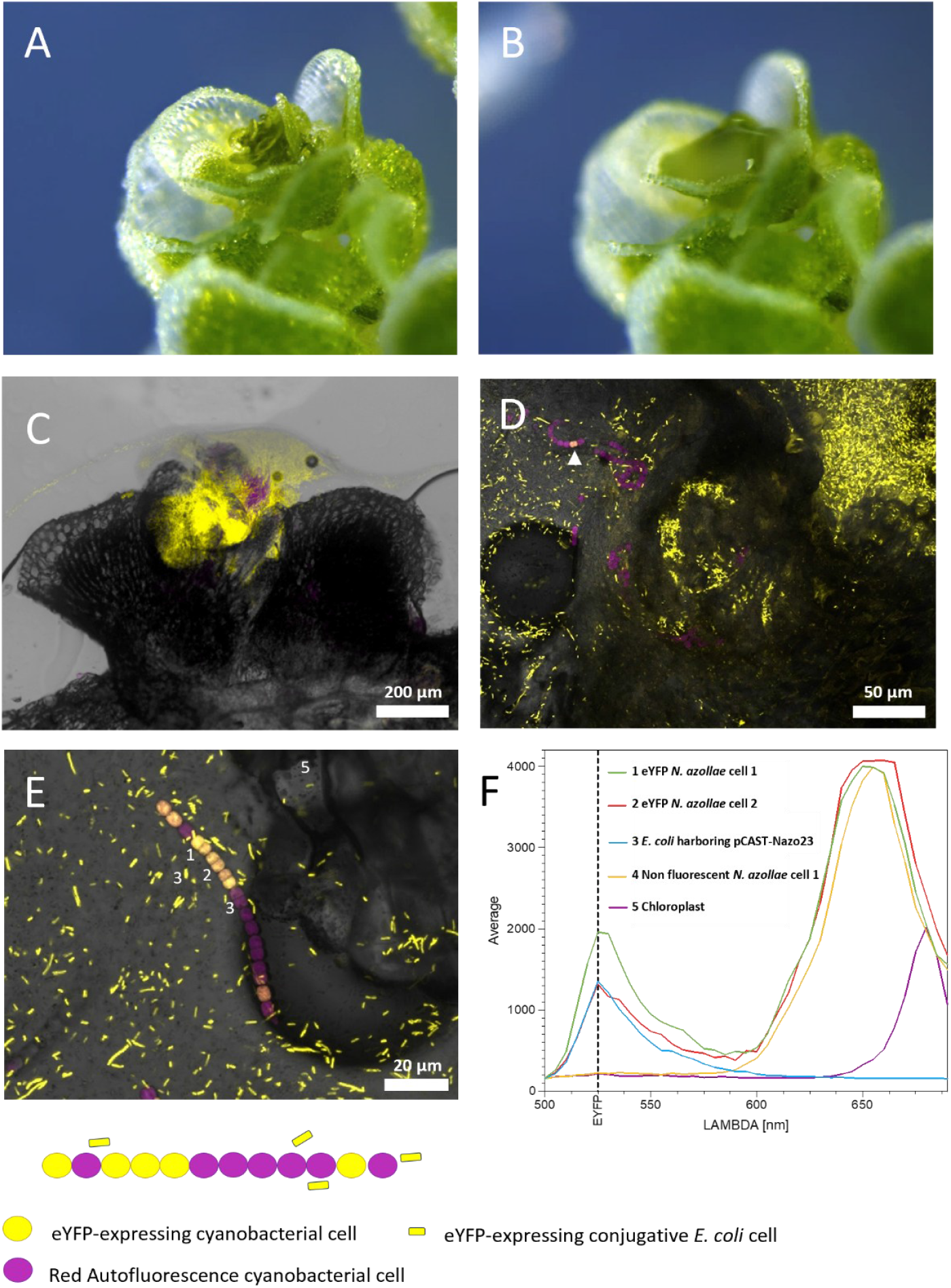
Conjugation of *Nostoc azollae in planta*. (A) Apex of *Azolla anzali* showing exposed meristem after treatment with 1 µg/mL BAP for 7 days, followed by 13 days without hormones. (B) Apex from “A” immediately after the application of concentrated conjugative *E. coli* suspension with microneedles. (C) Example of inoculated apex of *Azolla* observed by confocal microscopy 24 hours post-inoculation. (D) Evidence of a single conjugation event of *N. azollae* in *Azolla anzali* 5 days post-inoculation. Confocal images composed by three merged channels: BF: Brightfield, RAF: Red autofluorescence (in magenta) and eYFP channel. (E) Z stack detail of *N. azollae* filament showing 3-4 conjugation events. Average fluorescence emission spectra corresponding to the numered spots from a single focal panel of micrograph from E panel. Conjugation was performed with *E. coli* harboring pCAST-Nazo23. A scheme depicting the different cells observed, expressing or not expressing the eYFP, is also included.

To further characterize SAM structure and SANC properties, a staining procedure using the fluorescent DNA-binding dye 4′,6-diamidino-2-phenylindole (DAPI) was performed (Fig. S9). The results showed that it was possible to manually deliver a payload to the SANC and helped us to determine optimal imaging conditions for *in planta* confocal microscopy. Once the SAMs were exposed (Fig. 3A), the meristems were “massaged” using glass-made microneedles to facilitate application of a cream-consistency conjugative mix of *E. coli* cells (Fig. 3B, Fig. S3B). The “cream- like” consistency was needed to avoid the substantial hydrophobicity of *Azolla* tissue (28) and likely helped to increase contact between *E. coli* cells and SANC filaments by maintaining the *E. coli* payload in place. Following this procedure, the pL0 and pCAST-Nazo plasmids previously validated in *Azolla* juice and which drive eYFP expression from P_trc1O_ were used as cargo plasmids to trace *E. coli* location and successful inoculation (Fig. 3C), as well as the subsequent conjugation events (Fig. 3D-3E). After apex dissection and gentle squashing to allow confocal microscopy, eYFP fluorescence signals were detected within *N. azollae* cells (Fig. 3E) (see summary of experimental attempts and results in Table S3) and further confirmed by Lambda scanning (Fig. 3F). These results demonstrated successful plasmid transfer into the symbiont, *in situ*. During *in planta* experiments, *N. azollae* filaments with eYFP fluorescent cells frequently appeared “healthy”, without the fragmentation and pigment loss that were observed in *ex planta* conjugations (Fig. 2 and Fig. S5). Exconjugants were observed as early as 48 hours post-treatment, and could be detected also after 5 days, with no events found around 24 hours after mating (Table S3, Fig. S10A and Fig. 3). This is similar to conjugation detection times in *Anabaena*. After 48 hours, adjacent fluorescent cells within a filament were frequently observed, consistent with local proliferation of the exconjugant subpopulation, which is a key factor for potential spreading of a mutant (Fig. S10A).

Most *N. azollae* filaments with exconjugant cells were found in populations with characteristic features of the filaments in the SANC (narrow cells, short filaments without heterocysts), likely because they originated from the dissected SAM that was inoculated. This confirmed that this cyanobacterial population was successfully targeted (Fig. S10A). In a few instances, these SANC filaments were recorded clearly surrounding SAM structures including their large trichomes (Fig. S11; Movie S1). Additionally, exconjugant cells were occasionally observed entangled with heterocyst-harboring filaments that are typically found in leaf-pocket populations (Fig. S10B), which may represent exconjugants present in early populations forming within leaf pockets. This could reflect either (i) direct transfer from *E. coli* into *N. azollae* within the earliest developing pockets, or (ii) movement of exconjugant cyanobacteria from the SANC into adjacent compartments within the 48-h post conjugation. Distinguishing between these possibilities will require experimentation involving long-term survival of the treated host.

### Promoter characterization in *Nostoc azollae*

The suitability of promoters used to express each CAST-plasmid element (Fig. 4A) was assessed. Besides P_*trc1O*_ (driving eYFP in the CAST plasmid; Fig. 4B), we tested the synthetic P_J23119_ promoter (sgRNA expression; Fig. 4C), P_*glnA*_ (Cas12k and TnsBCD; Fig. 4D), P_*cpc560*_ (erythromycin [Em] resistance; Fig. 4E), and P_*psbA2L*_ (Fig. 4F) (Table S1). All promoters yielded observable conjugation events in *N. azollae* with measurable fluorescence (Fig. S12).

**Figure 4.**
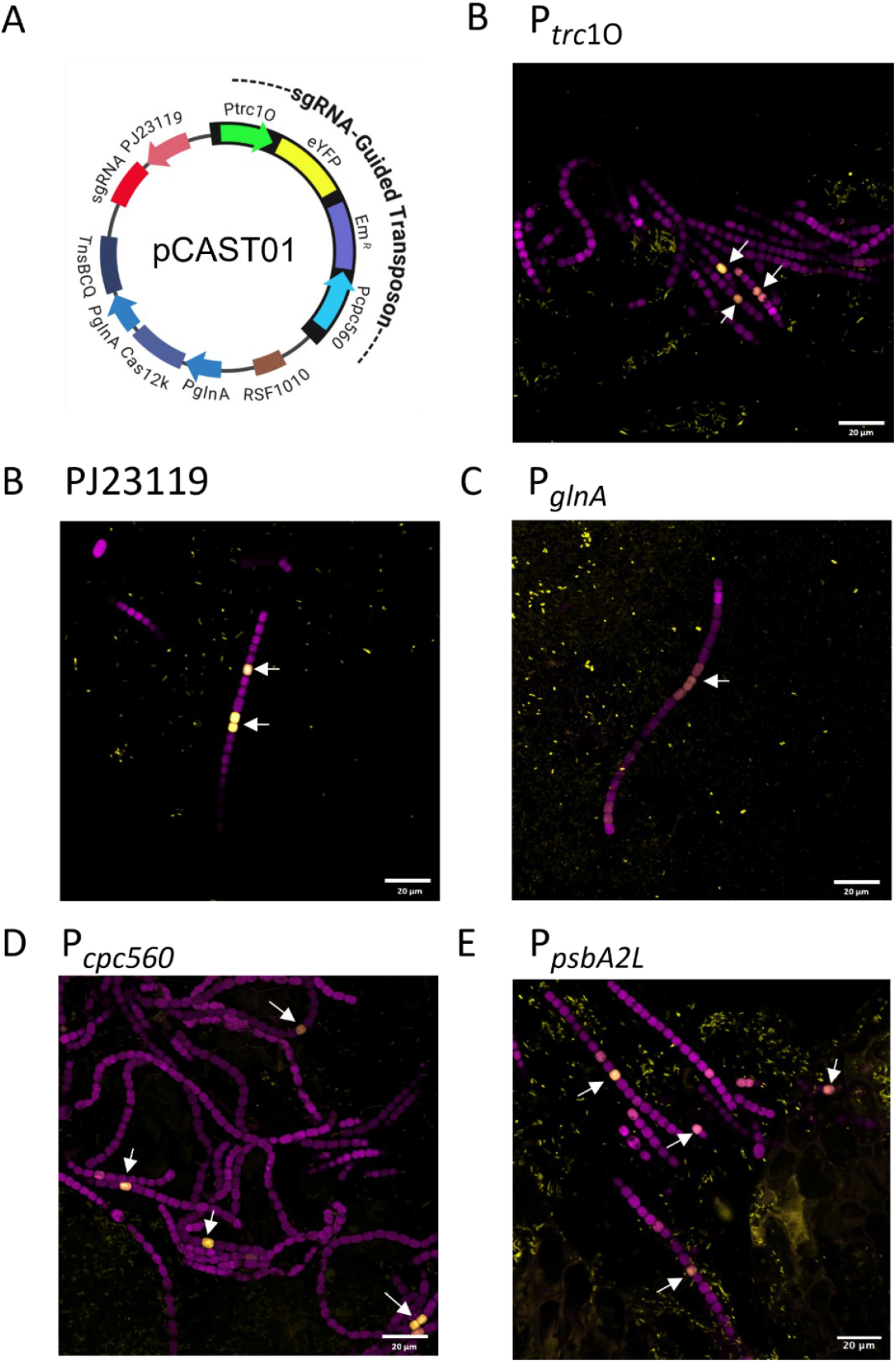
Testing of pCAST-related promoters in *Nostoc azollae in planta*. (A) Scheme of pCAST01 plasmid used to design CAST plasmids for *N. azollae* (pCAST-Nazo23 and pCAST- Nazo24) showing the promoters used to drive expression of each CAST system element (P_*psbA*2L_ is not shown in the scheme because it does not form part of pCAST01; however, it was also tested as an additional promoter option). Exconjugant cells from tests of plasmids with the following promoters: (B) P_*trc*1O_, (C) PJ23119, (D) P_*glnA*_, (E) P_*cpc560*_, and (E) P_*psbA2L*_. Some conjugation events are marked with a white arrow. Brightness and contrast were modified for improved visualization of exconjugant cells. All events were confirmed by Lambda scanning (Fig. S12).

Promoters were strongly expressed in *E. coli* except P_*cpc560*_ and P_*glnA*_, which showed weaker activity than in *N. azollae* (Fig. S12D–E). P_*cpc560*_, currently used for expression of Em resistance in pCAST-Nazo plasmids (Table S1), appears suitable for antibiotic selection: *N. azollae* exconjugants may tolerate Em while residual conjugative *E. coli* could be eliminated due to weak promoter activity. Importantly, Em has also been shown to be effective for generating cyanobiont- free *Azolla* strains (29). On the other hand, although most promoters produced numerous conjugation events, P_*glnA*_ yielded only a few detectable events. This low detection could be due to low heterologous P_*glnA*_ activity in the SANC population that may reflect its specific C/N regulatory status. In *Anabaena*, the complex P_*glnA*_ promoter is in part constitutive as well as being inducible by a high C/N cellular balance (30). In summary, detectable expression was observed for all pCAST- harbored promoters, suggesting that designing sgRNA-directed transposition could be a valuable tool.

Overall, *in planta* conjugation assays showed reproducible yet quantitatively variable outcomes across experiments (see summary of experimental attempts and results in Table S3). Despite standardized hormonal treatment and controlled growth conditions, the number of conjugation events varied markedly between apexes (from >200 events in some apexes to a few in others).

## Discussion

Metagenome sequencing and studies derived thereof have revealed the ubiquity and significance of the microbial contribution to the development and ecology of eukaryote species (31). *Azolla* ferns are no exception: without *N. azollae*, they grow poorly even in the presence of nitrogen nutrients, have a divergent community of associated microbes, do not undergo the transition to sexual reproduction and are not found in the natural environment (1, 29). Often, the microbial endosymbiont community cannot be cultured. To study host-microbe interactions, therefore, an important advance is the development of methods to manipulate genomes *in situ* as a prerequisite to identify genes mediating interactions. The present study demonstrates that *E. coli* may be used to achieve DNA-transfer to the filamentous cyanobacterial symbiont *in situ*, both, in cyanobacteria stem cells of the shoot apex and from early forming leaves where cyanobacterial filaments start differentiating N_2_-fixing heterocysts. This is the first evidence of DNA transfer to an obligate cyanobacterial symbiont. Our study further shows that the large RNA-guided transposition vectors originally designed for function in *Anabaena* (22) are efficiently transferred by *E. coli* into *N. azollae*, and that the promoters used to drive CAST-element expression in the plasmid are suitable for heterologous expression in *N. azollae in planta*.

The conjugation treatment appears to induce, however, a hypersensitive-like response in *Azolla*, preventing growth of the inoculated apexes (Fig. S13). Improving plant viability will be necessary to establish stable mutant populations. *In situ* genome engineering of bacterial communities using RNA-guided transposition in guts has been recently reported with *E. coli* (32), which is a natural and compatible endophyte in the gut. In contrast, in the *Azolla/Nostoc* plant/cyanobacterial symbiosis, whereas major advancements have been made in this study to accomplish *in situ* genome engineering, the challenge of overcoming the hurdle of compatibility with the host fern remains.

Despite the apparent hypersensitive response of *Azolla* against the conjugation procedure, the conjugative machinery encoded by pRL443 in *E. coli* worked successfully. Conjugative *pili* are assembled by type IV secretion systems (T4SS), which are known to be inactivated during host– pathogen interactions (Burdman et al. 2011). In the *Azolla* tissue, however, the conjugative T4SS of the *E. coli* donor was evidently not inactivated prior to plasmid transfer to *N. azollae*. This may be explained assuming an essential role of T4SS in the complex microbial community associated with *Azolla*, which includes Rhizobiales bacteria (29), as is the case in in *Bradyrhizobium*–legume symbioses (33). These observations align with those in the obligate *Burkholderia* leaf-nodule obligate symbionts of Rubiaceae plants, which, despite exhibiting pronounced genome erosion and strict vertical transmission, show evidence of horizontal gene transfer (34). In that system, plasmid conjugation has been proposed as a key driver of gene flow. Thus, although obligate symbioses might be perceived as evolutionarily static, the host may permit genetic transactions that enable continued genome plasticity in their microbial partners. The successful conjugation of *N. azollae* shown here indicates a long-term retention of the capability to receive DNA by conjugation despite the ∼100 million years of evolution within the *Azolla* host (19, 20), which may contribute to a genome plasticity in *N. azollae* that warrants further research.

## Materials and Methods

### *Azolla* fern and cyanobacterial cultures

Two different species of *Azolla* were used, *Azolla filiculoides* strain Galgenwaard (20) and *Azolla anzali* (17). *Azolla* sporophytes were cultivated in SAM (Standard *Azolla* Medium), a variation of IRRI2 medium (17), at 20ºC under a 16-hour light/8-hour dark cycle; light period (100 µmol m^−2^ s^−1^, LED-tube 4000K) was supplemented with Far-Red light from an APEXstrip Bundle 730 nm – 16W system (Crescience). The cyanobacterium *Anabaena* sp. strain PCC 7120 (hereafter *Anabaena*) was cultured photoautotrophically in BG11 or BG11_0_ (BG11 without NaNO_3_) media at 30°C, under constant light (30 µmol photons m^−2^ s^−1^, LED-tube 4000K) unless stated. BG11 medium was as described (35) with ferric ammonium citrate substituted by ferric citrate. Liquid cultures were maintained in Erlenmeyer flasks and shaken. For cyanobacterial solid cultures, 1% (w/v) agar (Bacto-Agar, Difco) was added. When required, growth medium was supplemented with the indicated concentrations of antibiotics.

### Plasmid constructions and cloning procedures

All plasmids generated in this work were developed using Golden Gate cloning and are summarized in Table S1. Level 0 plasmids were built with previous parts (22) such as P_glnA_ and PCR-adapting new individual components obtained from CyanoGate (24) or MoClo Plant level 0 plasmids (36). Level 1 plasmids were assembled by digestion with *Bsa*I and ligation. To obtain level T plasmids, level 1 inserts were assembled into pCAT.000 (conjugative and replicative), together with an end-linker (L) for CAST plasmids, by *Bpi*I digestion and ligation. The desired sgRNA was cloned with one *Lgu*I reaction either into Level 1 pAzU1.3 plasmid (22) or directly into the final pCAST empty plasmids named pCAST01 (Em^R^) and pCAST02 (SpSm^R^). Sequences of oligonucleotides used in this work are summarized in Table S2. Final pCAST vectors ready to clone any sgRNA by one *Lgu*I reaction will be made available upon request. Reactions for Golden Gate assembly of Level 1 and Level T plasmids and sgRNA oligonucleotide annealing are described in Supporting information. Restriction enzymes and T4-DNA ligase were from ThermoFisher. Plasmids were introduced in *E. coli* DH5α and HB101 by heat shock (37). Putative positive colonies were selected using appropriate antibiotics and blue–white screening (38). Positive colonies were confirmed either by PCR with Taq polymerase using specific primers or by plasmid isolation and restriction. All plasmid sequences were finally checked by nanopore sequencing (Plasmidsaurus).

### Triparental mating procedure for *Anabaena* and *Nostoc azollae* extracted from *Azolla* juice (*ex planta* conjugation)

Triparental mating was performed using *E. coli* ED8654 (harboring pRL443, providing the conjugation machinery) and *E. coli* HB101 (carrying the helper plasmid pRL623 and the cargo plasmid), following the procedure of Elhai and Wolk (1988)(39). Overnight precultures of both *E. coli* strains were used to initiate fresh cultures by inoculating 10 mL of Luria–Bertani (LB) medium (without antibiotics) with 250 µL in the case of ED8654 preculture and 350 µL for the HB101 preculture. Refreshed cultures were incubated for 2.5 h at 37 °C with shaking to reach exponential growth. Cells from donor and helper cultures were then transferred to round-bottom tubes and gently centrifuged at 1600 × *g* for 3 min at RT. Supernatants were removed, and pellets were resuspended in remaining LB medium and mixed by gently tapping the tubes. Donor and helper cells were incubated statically at 25 °C for 2–4 h and then mixed with filamentous cyanobacteria: an amount of cells containing 10 µg of chlorophyll *a* (Chl) for *Anabaena* and 20 µg Chl of *Azolla* juice containing *N. azollae* obtained as previously described (7) using BG11_0_ buffered with 10 mM TES (pH 7.5) as extraction media to prevent possible pH variations due to release of *Azolla* trichome acidic content. Chl was determined in methanolic extracts of the cells as described (40). The higher amount of *N. azollae* cells compared to *Anabaena* was used to compensate for potential underestimation of cyanobacterial Chl due to remanent plant debris Chl. The cyanobacteria–*E. coli* mixture was gently spread onto nitrocellulose filters (Immobilon, Millipore REF: HAFT08550) placed on plates containing a solidified mixture of 95% BG11 or BG11_0_ (for *N. azollae* conjugation experiments with *Anabaena* used as control) medium and 5% LB. After 24 h, filters were transferred to BG11 or BG11_0_ medium, respectively. For selection in *Anabaena*, filters were transferred after 24–48 h to plates of BG11 medium supplemented with appropriate antibiotics (25 µg/mL neomycin, 5 µg/mL spectinomycin sulfate and 5 µg/mL streptomycin sulfate, or 10 µg/mL erythromycin). In *N. azollae ex planta* conjugation experiments, conjugal filters were incubated in darkness during the first 16-24h of mating and then maintained under a 12-h light/12-h dark photoperiod; white-light intensity, 30 µmol photons m^−2^ s^−1^.

### Preparation of *Azolla* for *in planta* conjugation of the obligate symbiont *Nostoc azollae*

A detailed scheme of the “*in planta” Nostoc azollae* conjugation is shown in Fig. S3.A. Access to the genetic target in the Shoot Apical *Nostoc* Colony (SANC) was performed by exposing the Shoot Apical Meristems (SAMs) of *Azolla* to cytokinin (BAP; the cytokinin 6-benzylaminopurine) treatment. Fresh growing *A. filiculoides* strain Galgenwaard or *Azolla anzali* cultures were used with cultures of 3-finger plantlets in 50 mL of fresh medium as starting point. After 1 week, they were sub-cultured in fresh medium with and without 0.5 mg/L BAP (for *A. filiculoides*) or 1 mg/L BAP (for *A. anzali*). Hormone treatment was performed for 24 hours based on experiments to determine the optimal hormone treatment protocol (see Supp. Fig. S7A). To remove BAP, plants were rinsed by vigorous swirling for 5 seconds in distilled water and transferred to fresh hormone-free culture medium. After 10 days, fronds started developing open meristems and were used for *in situ* triparental conjugation.

### Preparation and inoculation of *E. coli* conjugative mix for *in planta* conjugation

Two approaches, “classical” and “short”, were used for the preparation of the *E. coli* conjugative mix as depicted in Table S3. After evaluation, the “short,” protocol was chosen as it made it possible to inoculate more plants within the same day. However, the “classical” protocol was followed by using the *E. coli* conjugative mix prepared as mentioned above for *Anabaena*. After the biparental incubation, the *E. coli* mix was plated on a LB plate and left to dry before use. The “short” protocol consisted of preparing the *E. coli* conjugative mix using directly the overnight *E. coli* cultures instead of a refreshed exponential phase *E. coli* culture, as this showed little difference in conjugation efficiency (41). After gentle washing with fresh LB medium to get rid of antibiotics, *E. coli* cultures were mixed and 200 µL drops of the mix were placed on LB plates and let them dry. They were then incubated for 3–4 hours on the plate to obtain a *E. coli* conjugative “cream.” Occasionally, an O/N incubation of the cream mix on the LB plate was also used to reduce processing time.

To inoculate the *Azolla* apex with the *E. coli* conjugative cream, two glass microneedles (Glass replacement 1.14 mm 3.5” [(WPI]) were built using the Next Generation Micropipette Puller P-1000 (Sutter instrument) with the following parameters: program 2 at ramp 494 (Fig. S3.B) keeping the LB plate to avoid over-drying of the *E. coli* mix drop. Conjugative *E. coli* “cream” was carefully placed in exposed shoot apical meristems that showed the presence of SANC. After gentle “massaging”, the frond with a total of 2-3 treated apexes was carefully placed into a well (of a 24- well plate) with 1 mL of liquid SAM medium. After several trials (Table S3), to simplify the protocol and maximize standardization, the following conditions were selected. For the first 24 hours, plates were incubated in the dark at 30 ºC, followed by an additional 2 days at 30 ºC under a 16-hour light/8-h dark cycle; white-light period, 30 µmol photons m^−2^ s^−1^. For longer incubation times, *Azolla* fronds were placed in fresh growth media and transferred to regular *Azolla* growth conditions (20 ºC under a 16-hour light/8-h dark cycle; light period, 100 µmol photons m^−2^ s^−1^ supplemented with Far-Red light).

### Confocal microscopy and monitoring of conjugation events

Cyanobacterial conjugation was monitored through eYFP detection by confocal microscopy. For *Anabaena* conjugation and *ex planta* conjugation with *N. azollae*, a scrape of cells from a conjugation filter was streaked on a plate of BG11_0_ medium, then cut out of the agar and covered with a coverslip for visualization. For screening of *in planta* conjugation, *Azolla* apexes were dissected under a Leica M205 C Stereo Microscope and gently squashed between two coverslips in the presence of 20 µl of 60% glycerol to avoid desiccation. Images were collected with an Olympus FLUOVIEW FV3000 confocal laser-scanning microscope equipped with a UPlanApo 40 × 1.5 NA oil immersion objective. The eYFP was excited using a 488-nm laser at 2% power, with irradiation from an argon ion laser. The fluorescence of eYFP was visualized with a window of 500−540 nm, and the autofluorescence from the natural pigments of cyanobacterial cells was collected using a window of 660−700 nm. ImageJ-FIJI (42)/Olympus software was used to process images.

The conjugation success was followed using confocal microscopy at the selected time points after conjugation. To quantify conjugation events in *N. azollae in planta* experiments, consecutive eYFP expressing cells were considered as an individual conjugation event followed cell division. Lambda scans, routinely performed to discern eYFP signal from cyanobacterial autofluorescence signal, were recorded over the 500–700 nm spectral window following excitation with a 488-nm laser at 2% power. The average fluorescence intensity within the region of interest was plotted as a function of wavelength.

## Acknowledgments

Work was supported by the Gordon and Betty Moore Foundation grant no. 9355. We thank Alicia Orea for help with confocal microscopy, Raquel González for technical support and maintenance of the *Azolla* and cyanobacterial cultures, and José Manuel Pardo and Natalia Raddatz for providing the microneedle builder system and technical assistance with its use.

## Author Contributions

CSB designed and performed experimental work and interpreted data; CSB and HS set up conditions for preliminary attempts of *N. azollae* conjugation; EG and HS provided hormone treatment methodology and *Azolla* culture handling protocols; EF, PL, SNB, HS raised funds and supervised work; CSB, HS and EF drafted the manuscript; all authors provided input on the content of the manuscript and the figures.

