## Supplementary information for "Genetic access to an obligate cyanobacterial endosymbiont within the shoot meristem of fern hosts"

**This PDF file includes:**

Supporting information
Figures S1 to S13
Legend for Movie S1

**Other supporting materials for this manuscript include the following:**

Tables S1 to S3
Movie S1

**Supporting information**

**Reactions for Golden Gate assembly of Level 1 and Level T plasmids and sgRNA oligonucleotide annealing**

1. **Golden Gate reaction for Level 1 and Level T assembly.**

| **Part Type (insert/backbone)** | **DNA Amount (fmol)** | **Vol. DNA in Reaction (µL)[a]** |
|---|---|---|
| backbone | 20 | X |
| Insert (s) | 40 | X |
| | Buffer G 10X | 2.0 |
| | ATP 10 mM | 2.0 |
| | BsaI/BpiI[b] | 1.0 |
| | T4 ligase | 0.7 |
| | MQ (to 20 µL) | X |
| | **TOTAL (µL)** | 20.00 |

**[a] Volume of DNA in reaction taking into account the total Plasmid/PCR Length (bp) and the concentration of DNA (ng/µL)**
**[b] *Bsa*I enzyme was used for Level 1 and *Bpi*I for Level T assemblies.**

Thermocycler program:

| **Temperature (ºC)** | **Time (min)** | |
|---|---|---|
| 37 | 2 | 30-50 cycles * |
| 16 | 3 | |
| 37 | 5 | |
| 65 (for *Bsa*I) /80 (for *Bpi*I) | 10 | |

*** Increased cycling (50 cycles) was recommended for assembly of more than 5 parts or assemblies showing difficulties.**

2. **Reaction for sgRNA oligonucleotide annealing**

| | |
|---|---|
| F primer (500 μM) | 1 μL |
| R primer (500 μM) | 1 μL |
| T4 ligase buffer (x10) | 2 μL |
| Milli Q | 19 μL |
| **TOTAL** | 22 μL |

Thermocycler program: (95ºC for 5 min and cool down until 4ºC at a ramp of 5ºC/min)

3. **Golden Gate reaction to introduce sgRNA into Level T (pCASTX backbone) or Level 1 (pAzU1.3) plasmids**

| | |
|---|---|
| Buffer G 10X | 2 μL |
| ATP 10 mM | 2 μL |
| Vector (pCAST or pAzU1.3) | 100 ng (X μL) |
| sgRNA (hybridization reaction) | 300 ng (X μL) |
| LguI | 1 μL |
| T4 ligase HC (30 U/μl) | 0.4 μL |
| MilliQ | X μL |
| **TOTAL** | 20 μL |

Thermocycler program:

<table>
<tr><th>Temperature (ºC)</th><th>Time (min)</th><th></th></tr>
<tr><td>37</td><td>5</td><td rowspan="2">20 cycles</td></tr>
<tr><td>16</td><td>5</td></tr>
<tr><td>37</td><td>10</td><td></td></tr>
<tr><td>65</td><td>10</td><td></td></tr>
</table>

**Figures**

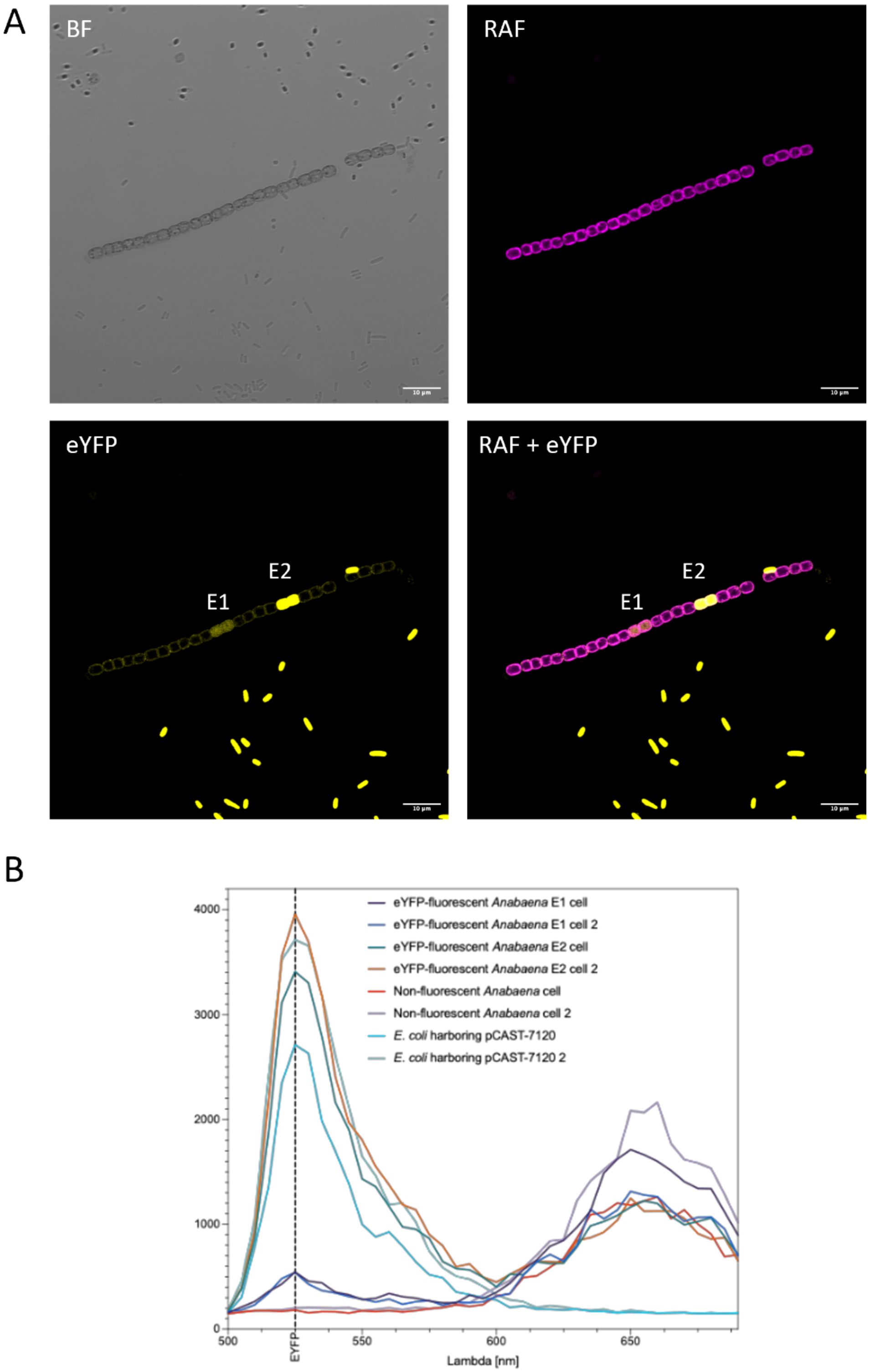


**Fig. S1.** Differential eYFP fluorescence intensity in two independent conjugation events and corresponding lambda scans. (A) Bright-field (BF), Red Autofluroescence (RAF, in magenta), eYFP, and merged RAF+eYFP images showing two conjugation events (E1 and E2) in *Anabaena* conjugation with pL0. In both events, conjugation-mediated DNA transfer resulted in detectable fluorescence likely followed by cell division; however, E2 exhibits higher eYFP signal compared to E1. Scale bar, 10 µm. (B) Corresponding lambda scans showing the fluorescence intensity of E1 and E2 events as well as controls in duplicate. Brightness and contrast adjusted to improve visibility.

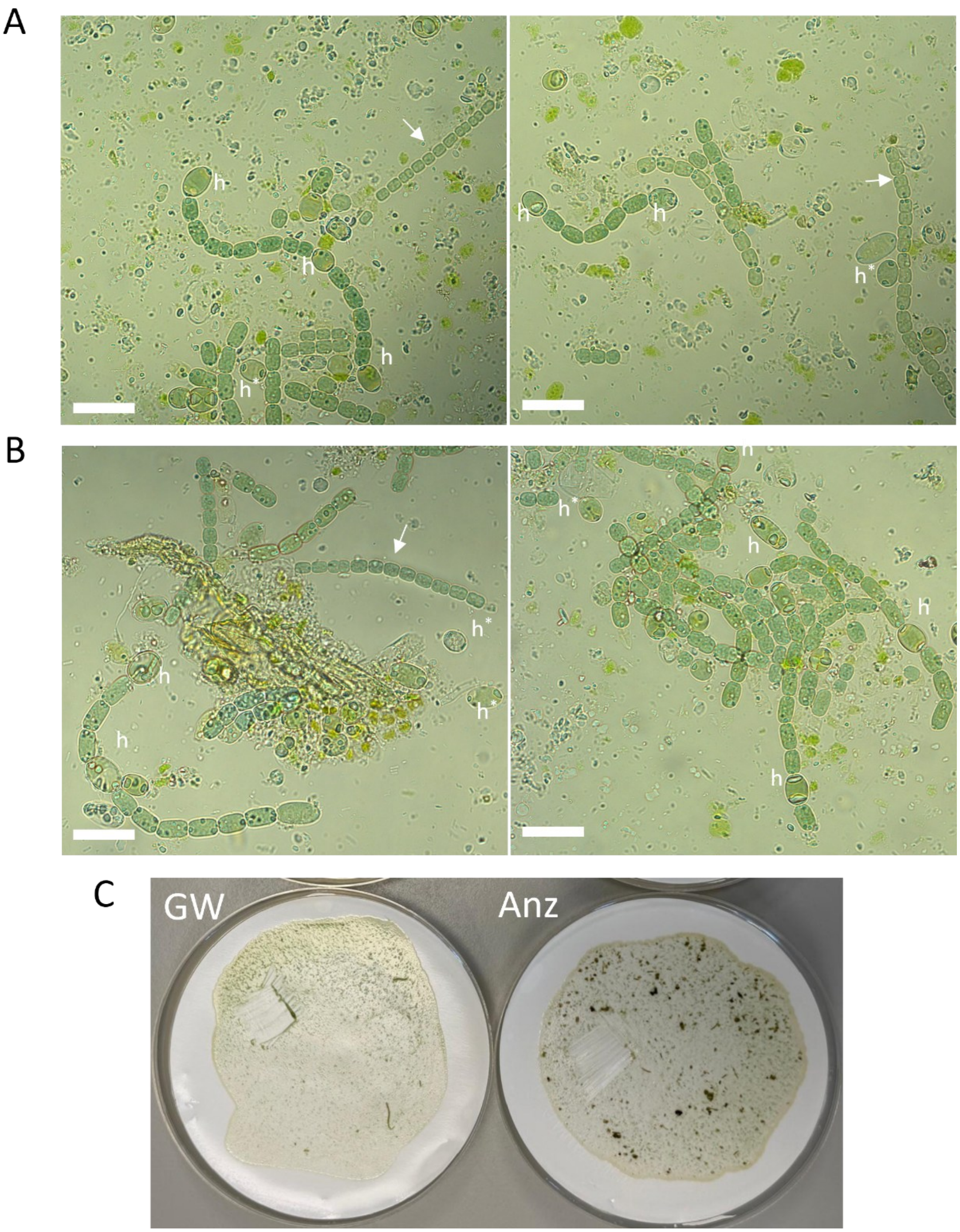


**Fig. S2.** Brightfield micrographs of *Nostoc azollae* filaments extracted following the method described in Sarasa-Buisán et al. (2025), with the addition of 10 mM TES (pH 7.5), and examples of conjugation plates. (A) Filaments isolated from *Azolla filiculoides*, strain Galgenwaard. (B) Filaments isolated from *Azolla anzali*. White arrows mark putative meristematic-like filaments/hormogonia composed of small-celled filaments lacking heterocysts. Some heterocysts are indicated (h) in filaments with larger cells. Note the presence of single heterocysts (h*), likely released from filaments indicating filament fragmentation. Scale bar, 20 µm. Micrographs were taken with a Leica DMRE microscope (C) Example of conjugation filters 48 hours post-mating of *N. azollae* filaments from juice extracted from *Azolla filiculoides*, strain Galgenwaard (GW) and from *Azolla anzali* (Anz). The scrap used for visualization of events in confocal microscopy is seen at the left hand of each filter. Note the presence of plant tissue debris in (A) and (B).

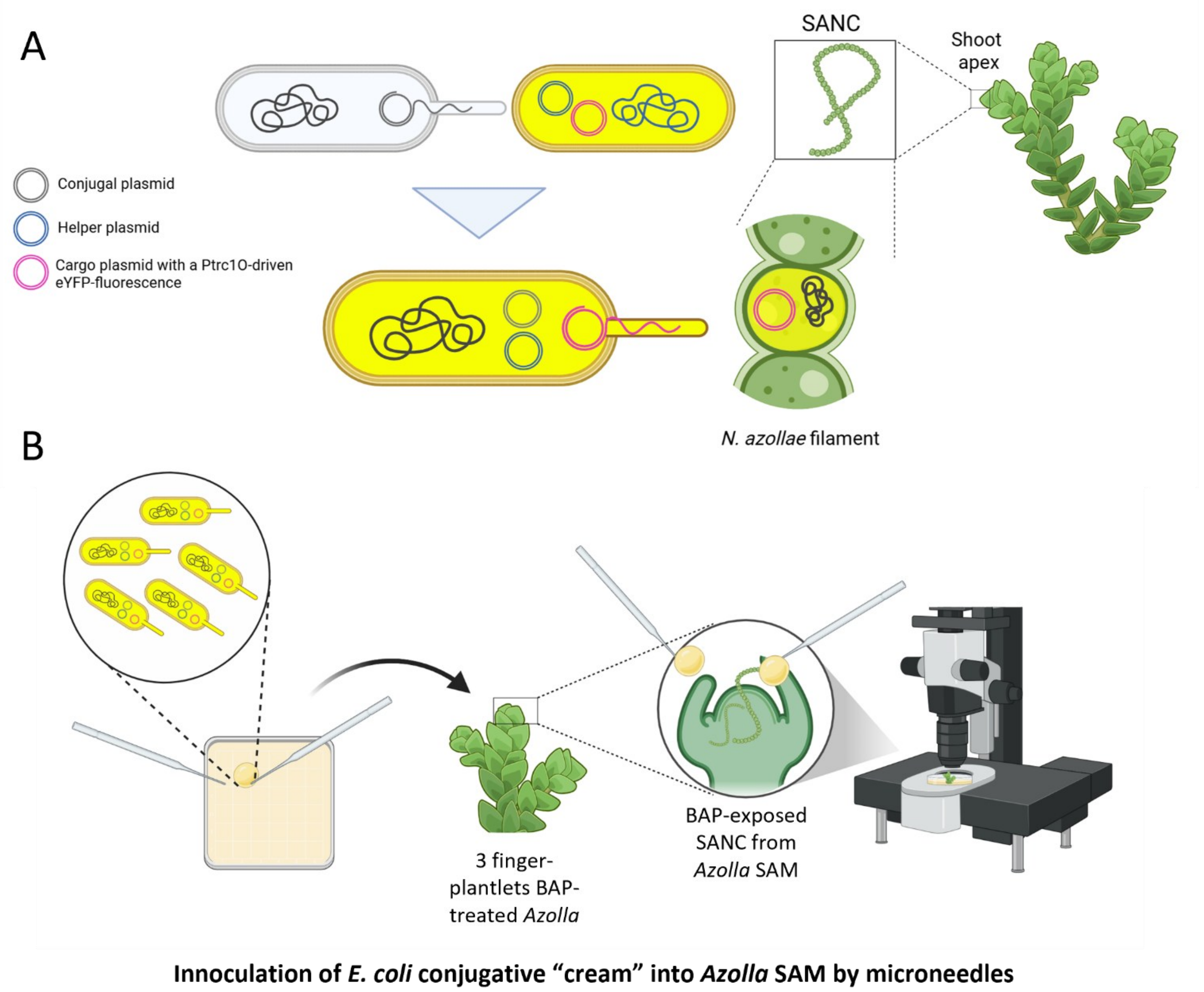


**Fig. S3.** (A) Scheme of *E. coli* triparental mating protocol developed for *N. azollae*. *E. coli/N. azollae* cells showing yellow cytoplasm represent eYFP fluorescence under the confocal microscope. (B) Protocol for *in planta* inoculation of conjugative *E. coli* cells into BAP-exposed SANC (Shoot apical *Nostoc* Colony) from *Azolla* SAM (Shoot Apical Meristem). Created with BioRender.com.

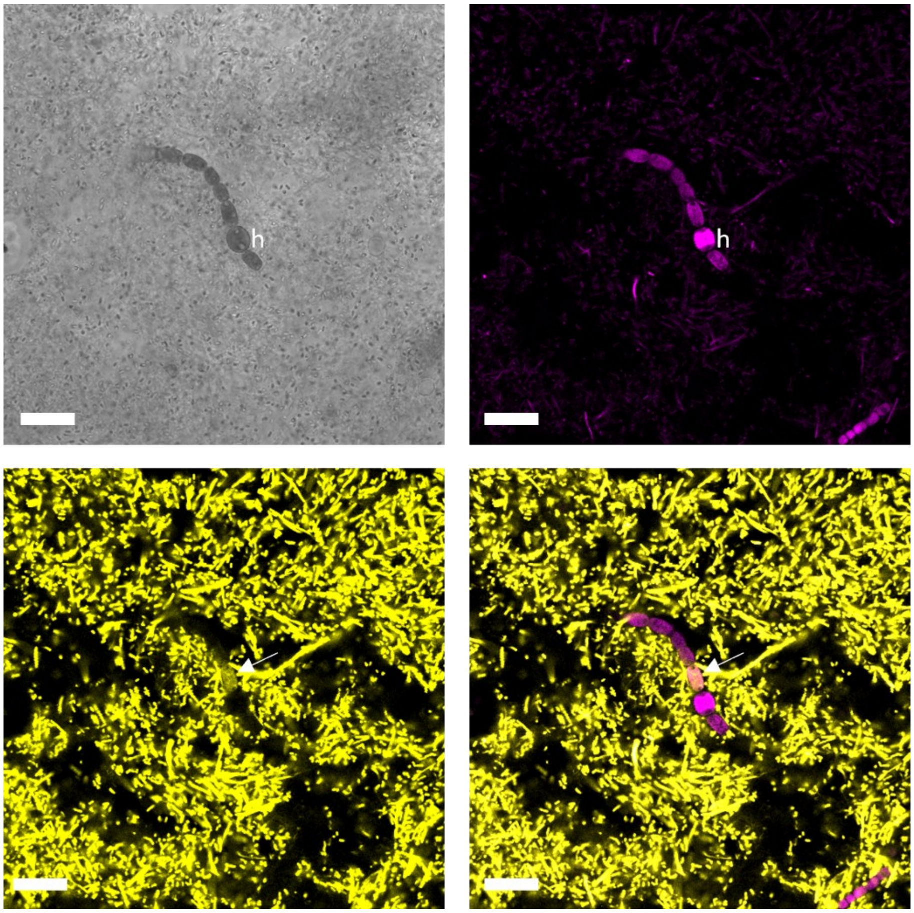


**Fig. S4.** Confocal microscopy of a conjugation event of *N. azollae* with pCAST-NazoT24 carrying the sgRNA directed to the *narB* pseudogene (used as a neutral platform). *N. azollae* cells were extracted from *Azolla filiculoides* Galgenwaard, and conjugation was performed and analyzed as described in Materials and Methods. Four detection channels are shown: BF (brightfield), RAF (red autofluorescence, shown in magenta), eYFP, and merged eYFP + RAF. Note the presence of a heterocyst (h), close to the eYFP-expressing cell indicated with a white arrow. Heterocysts show the higher RAF compared to vegetative cells characteristic in *N. azollae*. Scale bars, 20 µm. Brightness and contrast adjusted to improve visibility.

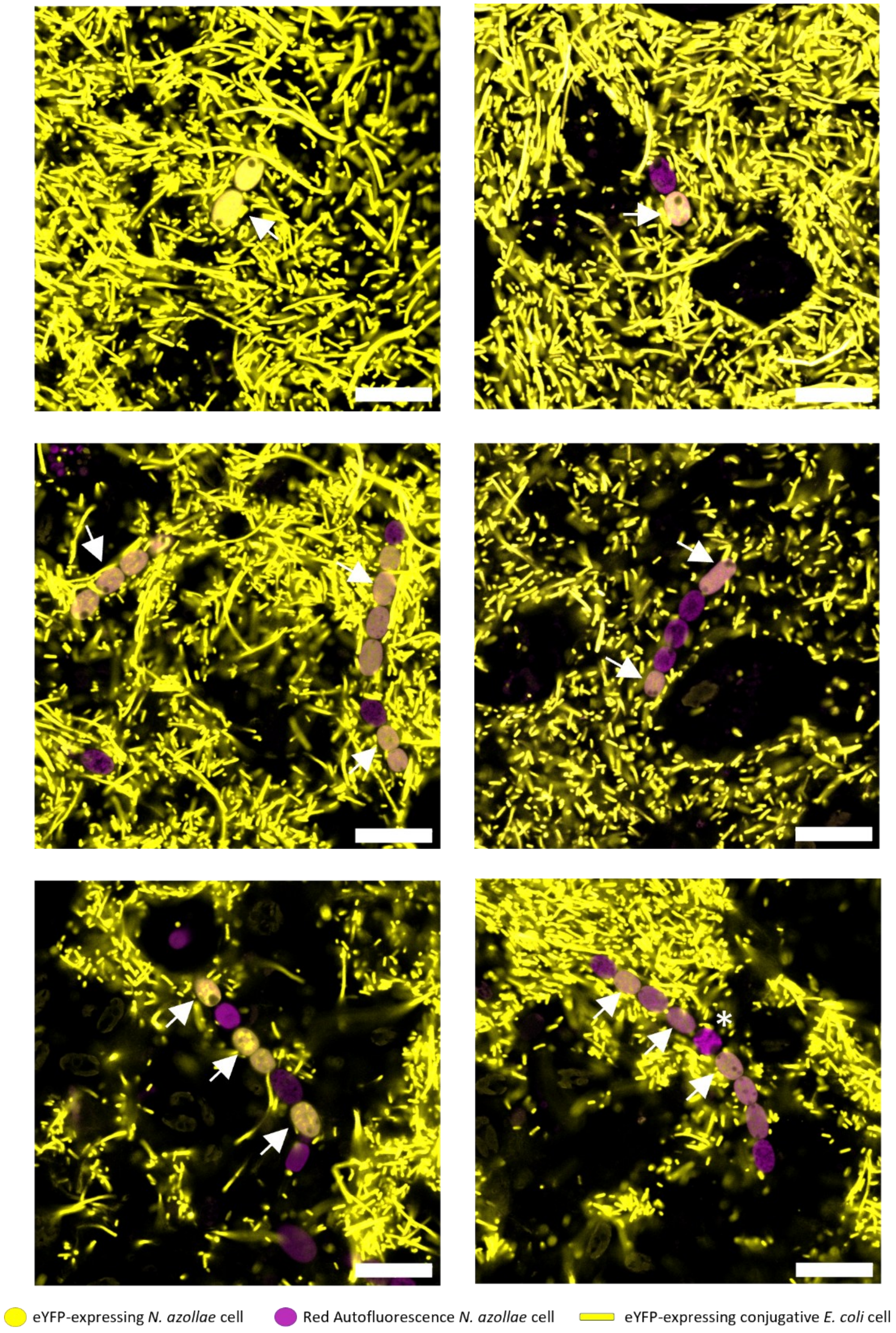


**Fig. S5.** Confocal microscopy of several representative configurations of conjugation events detected in extracted *N. azollae* filaments conjugated with pCAST-NazoT23 carrying a sgRNA directed to the *nifK*. *N. azollae* cells were extracted from *A. filiculoides* Galgenwaard, and conjugation was performed and analyzed as described in Materials and Methods. A merged eYFP + RAF channel is shown. Example of eYFP-expressing cells are indicated with white arrows, and a heterocyst is marked with an asterisk (*). Similarly detected conjugation events were also observed in the case of conjugation with pCAST-NazoT24. Scale bars, 20 µm. Images are representative of conjugation events observed in at least 3 independent experiments. A scheme depicting the different cells observed, expressing or not expressing the eYFP, is also included.

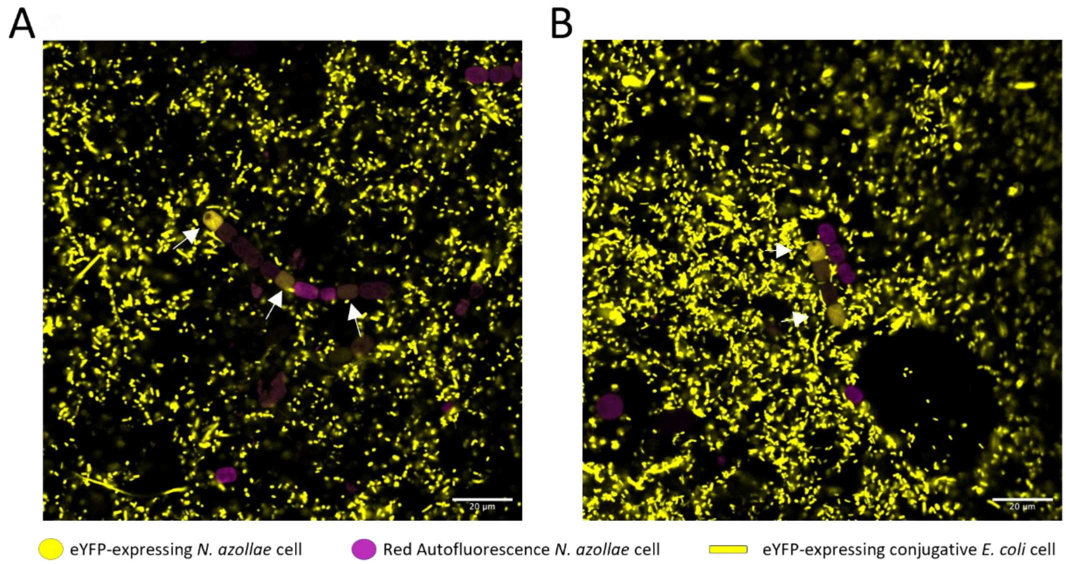


**Fig. S6.** Confocal microscopy of conjugation events detected in *N. azollae* filaments extracted from *Azolla anzali* with (A) pL0 and (B) pCAST-Nazo24. Conjugation was performed and analyzed as described in Materials and Methods. A merged eYFP + RAF channel is shown. Example of eYFP-expressing cells are indicated with white arrows. Images are representative of conjugation events observed in at least 3 independent experiments.

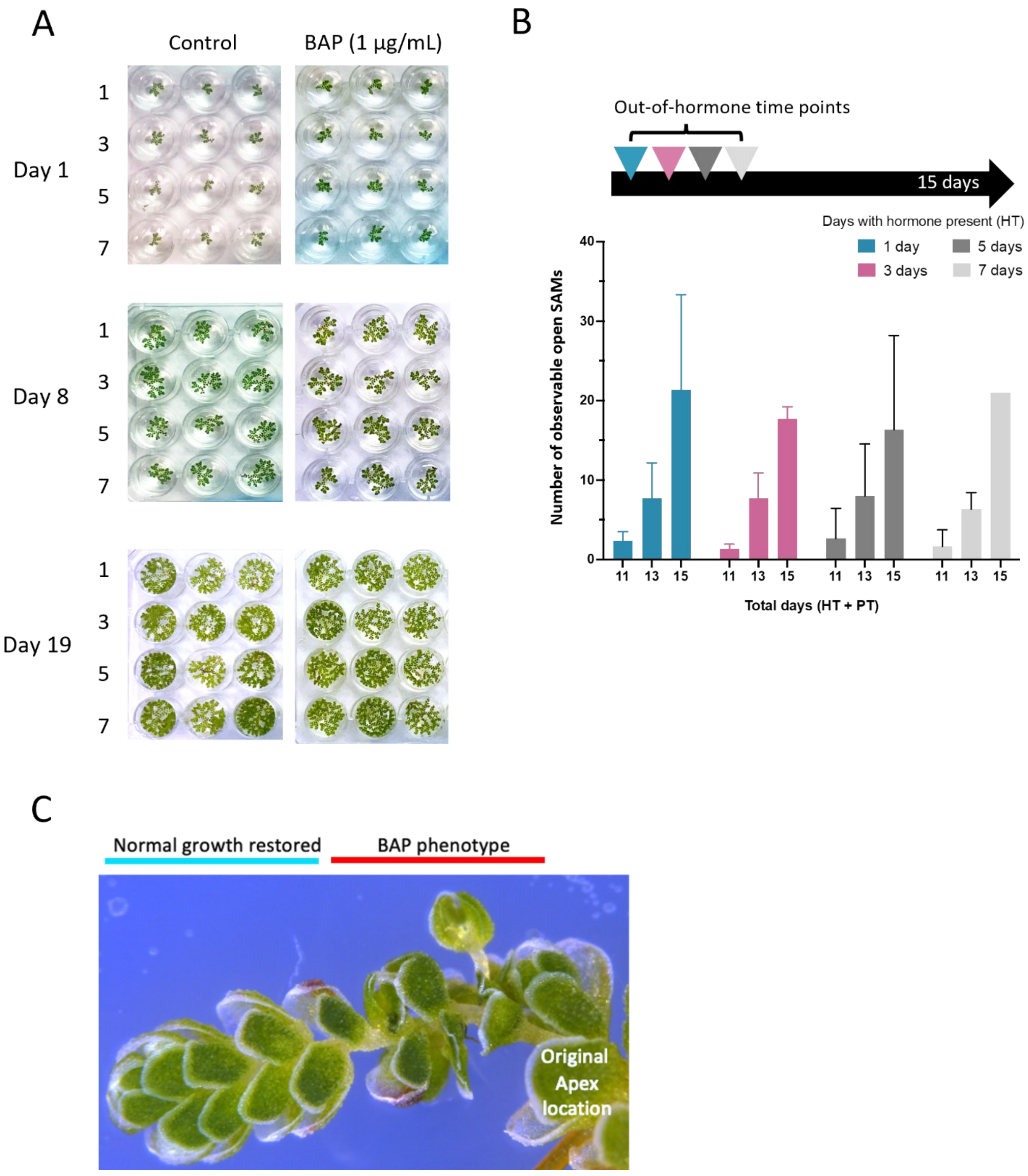


**Fig. S7.** Shoot Apical Meristem (SAM) exposure by cytokinin treatment. (A) Growth after incubation for several days in absence and presence of 1 µg/mL BAP. (B) Exposed SAMs counts during hormone treatment experiment. Hormone exposure was performed for 1, 3, 5, or 7 days, and phenotype was followed for a total of 15 days. Total number of days: HT (days of hormone exposure) + PT (days post-hormone). (C) Normal growth restored by outgrowth from BAP-phenotype apex. Original open apex is depicted.

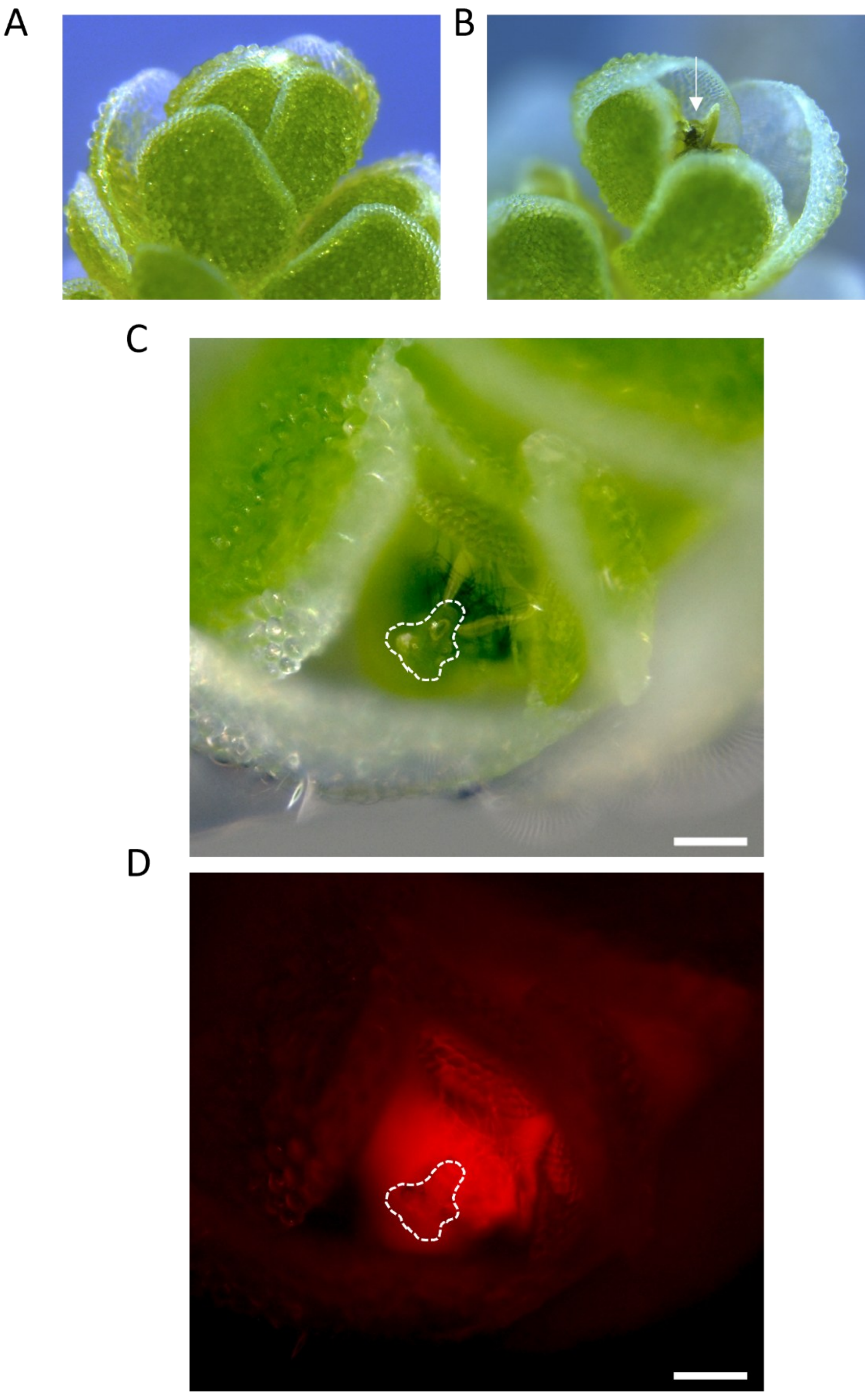


**Fig. S8.** Exposure of Shoot Apical *Nostoc* Colony (SANC) filaments from Shoot Apical Meristem (SAM) of *Azolla* by cytokinin treatment. Apex of *Azolla anzalli* before (A) and after (B,C,D) cytokinin treatment with 1 µg/mL BAP. Exposed SANC within SAM is marked with a white arrow in "B". (C) Close-up detail of SANC filaments surrounding SAM (delimited by a dashed white line) in brightfield. (D) Red autofluorescence image of "C" panel showing autofluorescence of cyanobacterial pigments localized surrounding the SAM. Scale bar 100 µm. BAP treatment was performed for 1 day, with the hormone phenotype observed from day 10 post hormone treatment.

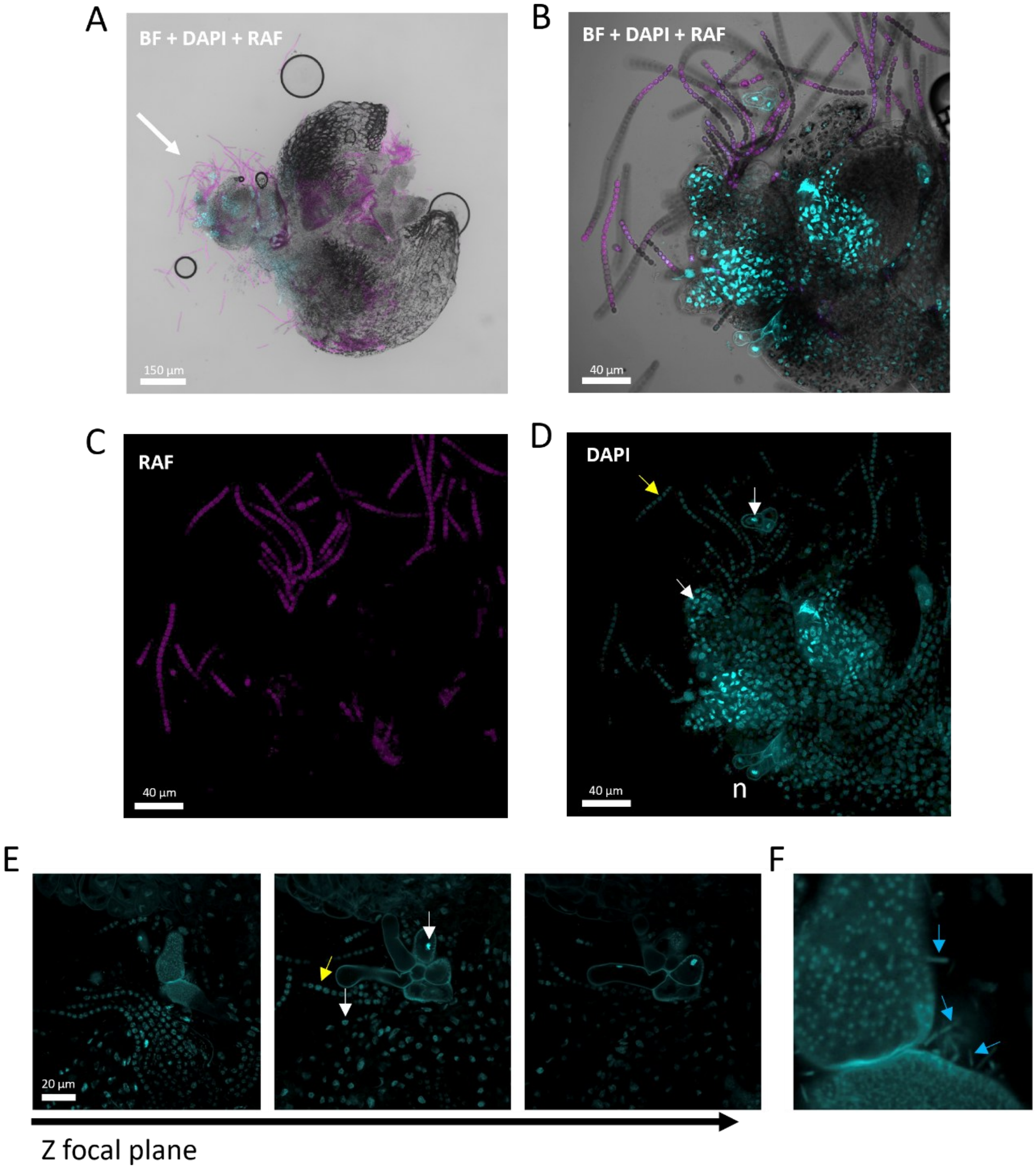


**Fig. S9.** DAPI labelling of a cytokinin-exposed Shoot Apical Meristem (SAM). *Azolla* apexes with exposed SAMs were isolated and placed on $BG11_0$ agar plates, followed by the addition of a 5 µL drop of a 10 µg/mL solution of the fluorescent DNA-binding dye 4′,6-diamidino-2-phenylindole (DAPI) prepared in Milli-Q water. After 15 min of incubation, apices were mounted in a 10 µL drop of 60% glycerol on a microscope slide and visualized by confocal microscopy using 405 nm excitation. (A) Micrograph of the whole apex at 10x magnification; a white arrow indicates the treated SAM. (B,C,D) Close-up of the DAPI-stained SAM harboring the Shoot Apical *Nostoc* Colony visualized in (B) Brightfield + DAPI + RAF (Red Autofluorescence) merged channels, (C) RAF channel and (D) DAPI channel. *Azolla* meristem was exposed by treatment with 1 µg/mL BAP for 7 days, followed by 13 days without hormone. Staining of the SAM was successful, specifically staining both nuclei of *Azolla* plant cells (including those in trichomes) (white arrow in panels (D,E)) and the DNA-containing cytoplasm ("bacterial nucleoid") of *N. azollae* (an example indicated by a yellow arrow in panels (D,E), confirming that the SANC target was successfully reached and visualized. *N. azollae* filaments close to the SAM appeared as small-celled filaments lacking heterocysts, and a two-lobed trichome that likely corresponds to the structure that attracts cyanobacteria to nascent leaves in the meristem is indicated by a white arrow. Notably, DAPI staining also revealed the location of putative heterotrophic bacteria associated with *Azolla* trichomes. (E) Three consecutive images that correspond to the same area but at different Z focal planes, showing that the stained "bacteria-like structures" are peripheral to the trichome. *Azolla* cells nuclei white arrow) and "bacterial nucleoid" of N. azollae (yellow arrow) are shown. (F) Close-up of panel (E) that highlights stained bacteria-like cells detached from trichomes (blue arrows).

A

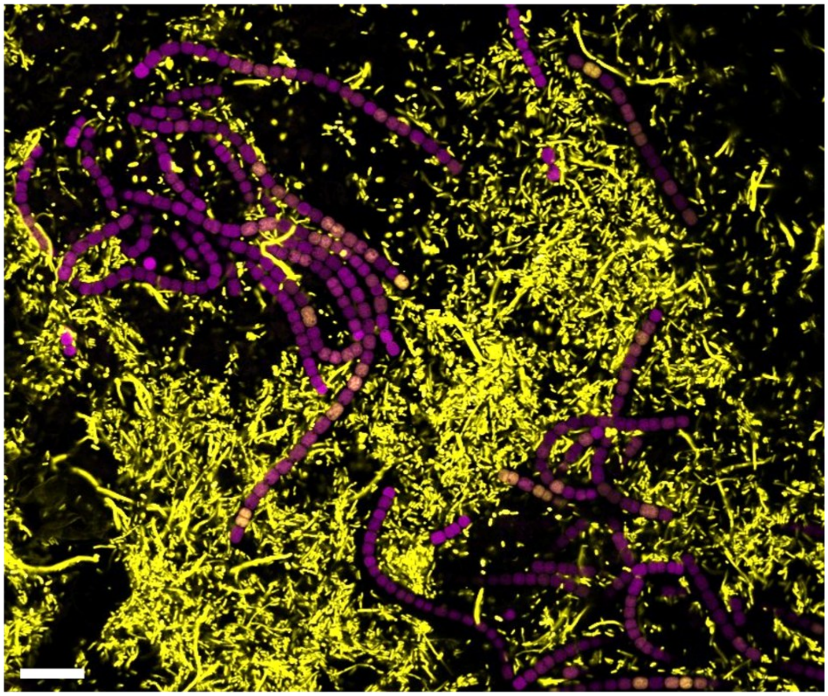

B

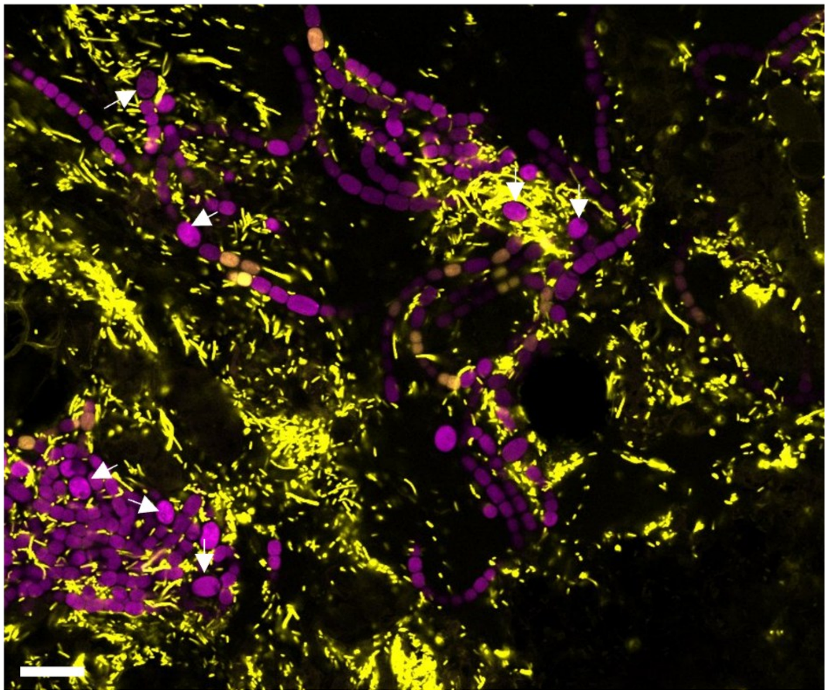

**Fig. S10.** *In planta* early conjugation events detected in two different *N. azollae* populations. Confocal microscopy image showing eYFP fluorescence and Red Autofluorescence (RAF, in magenta) of *N. azollae* filaments from (A) a population of SANC-like appearance, and (B) a population of leaf-pocket appearance. Note the presence of heterocysts (some depicted by white arrows) and more aggregated filaments with larger cells in the leaf-pocket appearance population (panel B). Cells were conjugated with the control plasmid pL0, which carries the $P_{trc10}$ promoter driving eYFP expression. The image was taken 48 hours after conjugation. *E. coli* donor cells expressing eYFP are also visible. Scale bar, 20 µm.

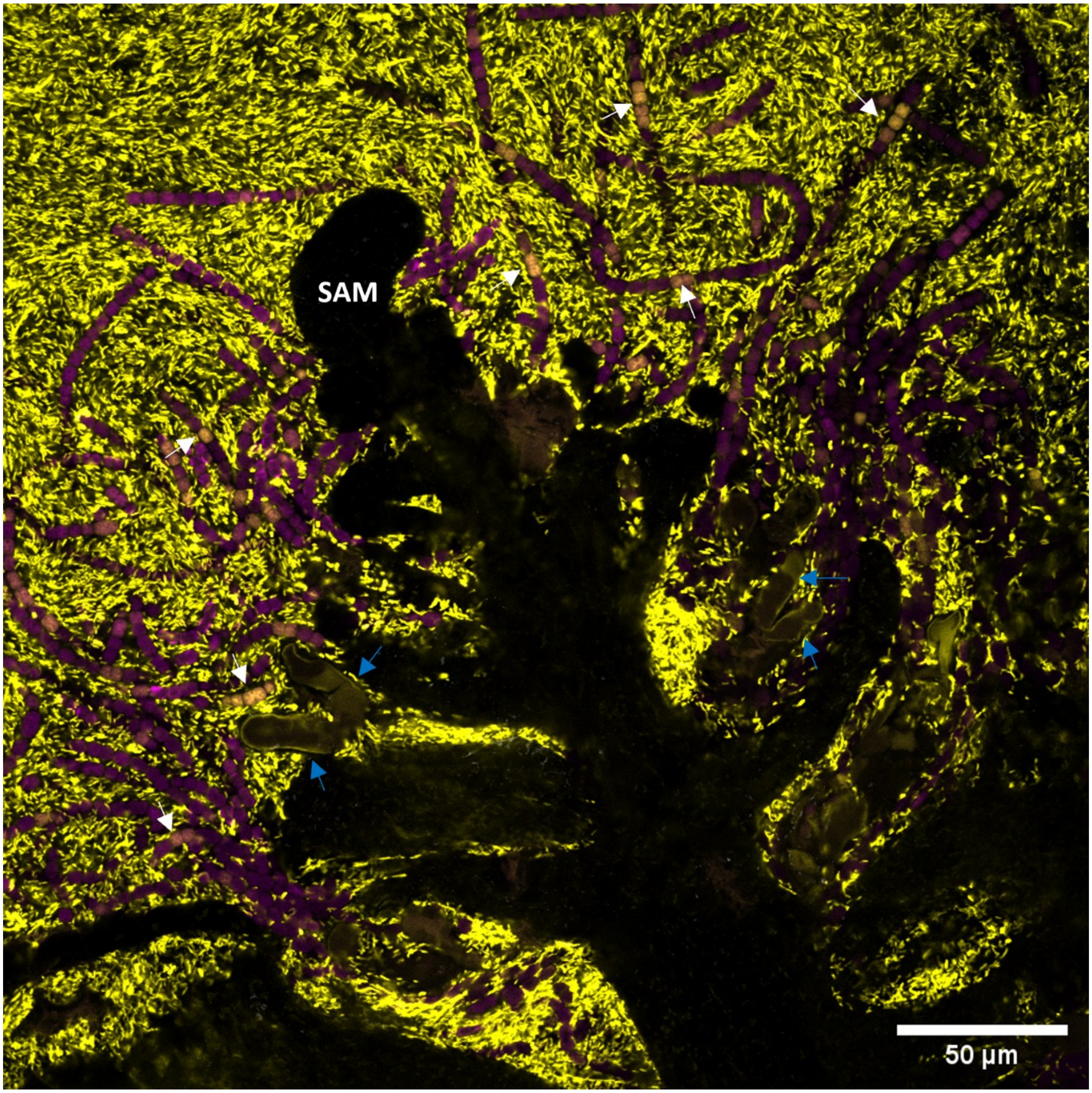


**Fig. S11.** *Nostoc azollae* filaments harboring conjugation events in *Azolla anzali* SANC. The image corresponds to a screenshot of a focal plane of Movie S1, in which many additional conjugation events can be observed surrounding the SANC structure. Confocal microscopy image combining brightfield, eYFP fluorescence, and red autofluorescence (RAF, shown in magenta) of *N. azollae* filaments located close to the indicated SAM-like structure. Numerous conjugation events are visible, some of which are indicated by white arrows. Large trichomes are indicated with blue arrows. Note also the absence of heterocysts. Cells were conjugated with the control plasmid pL0, which carries the $P_{trc10}$ promoter driving eYFP expression. The image was acquired 48 hours after conjugation. *E. coli* donor cells expressing eYFP are also visible.

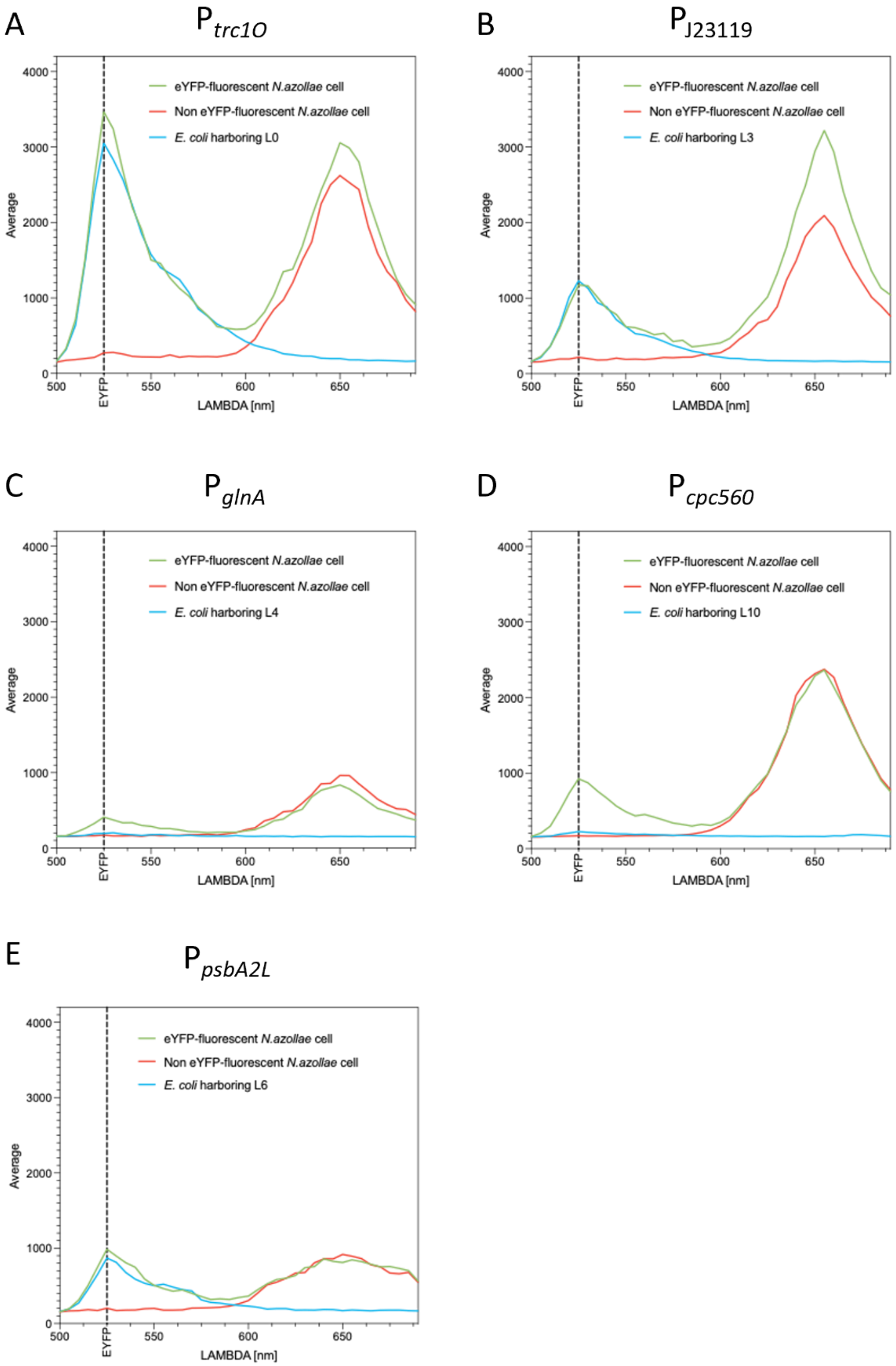


**Fig. S12.** Lambda scans of exconjugant cells from events of conjugation with plasmids testing the strength of different promoters included in pCAST plasmid. (A) $P_{trc1O}$; (B) $P_{J23119}$; (C) $P_{glnA}$; (D) $P_{cpc560}$; (E) $P_{psbA2L}$.

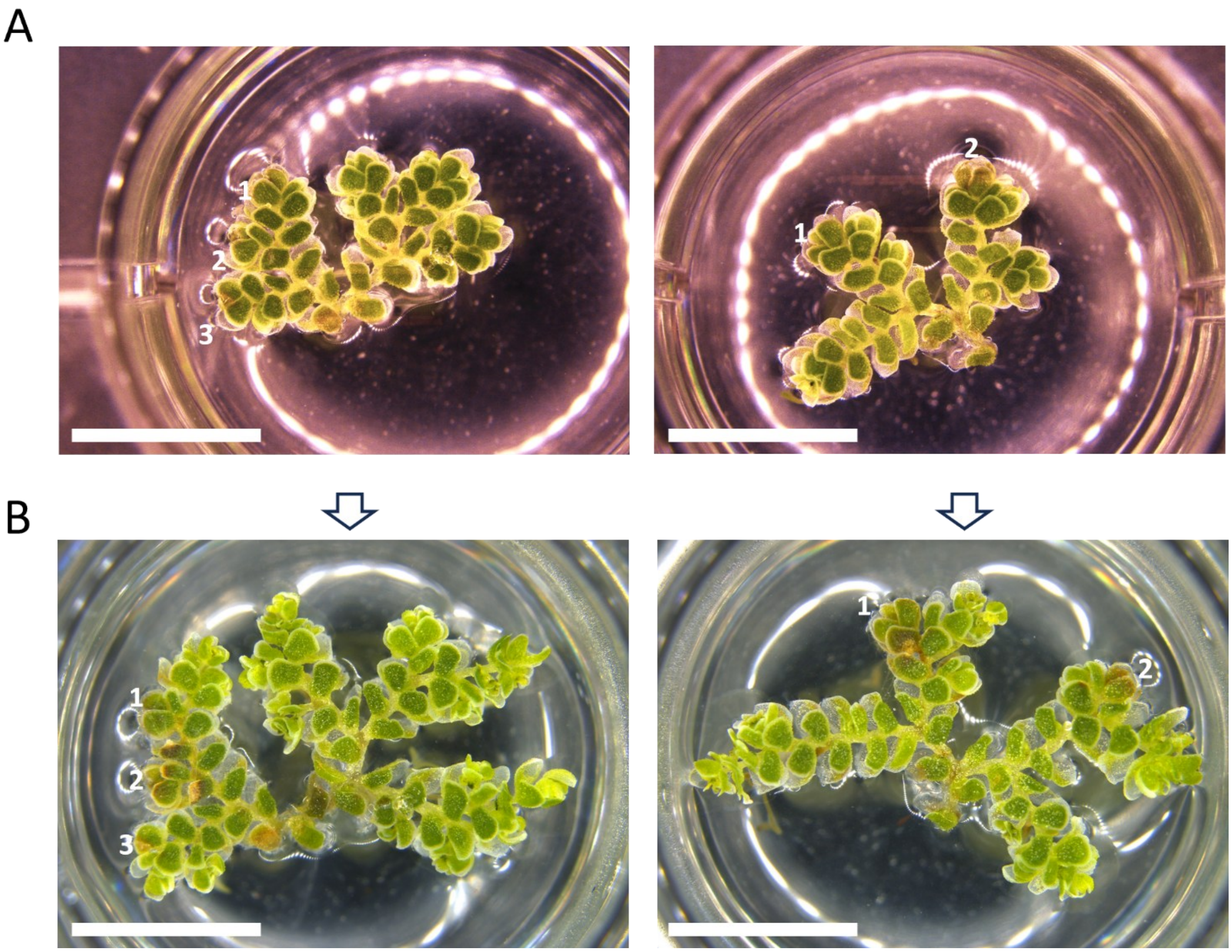


**Fig. S13.** Outgrowth monitoring of *Azolla anzali* apexes subjected to *in planta* conjugation. (A) Two examples of *Azolla* fronds with treated apexes (numbered in white) 48 hours post-inoculum. (B) Same fronds as panel (A) 10 days post-inoculum with corresponding numbered treated apexes. Note the stop of growth of treated apexes compared to non-treated apexes. Scale bar: 5 mm

**Table S1 (separate file).** List of Golden Gate-cloning plasmids used in this study.

**Table S2 (separate file).** List of oligonucleotides used in work.

**Table S3 (separate file).** Summary of results of the *in planta N. azollae* conjugation experiments.

**Movie S1 (separate file).** Movie showing animation through successive focal planes of *Nostoc azollae* filaments harboring conjugation events surrounding *Azolla anzali* SAM-like structure. Confocal microscopy image combining brightfield, eYFP fluorescence, and red autofluorescence (RAF, shown in magenta). Please see a representative screenshot in Fig. S11 highlighting the SAM-like structure, conjugation events and large trichomes. Note also the absence of heterocysts. Cells were conjugated with the control plasmid pL0, which carries the $P_{trc10}$ promoter driving eYFP expression. The images were acquired 48 hours after conjugation. *E. coli* donor cells expressing eYFP are also visible.
